# LUXendins reveal endogenous glucagon-like peptide-1 receptor distribution and dynamics

**DOI:** 10.1101/557132

**Authors:** Julia Ast, Anastasia Arvaniti, Nicholas H.F. Fine, Daniela Nasteska, Fiona B. Ashford, Zania Stamataki, Zsombor Koszegi, Andrea Bacon, Stefan Trapp, Ben J. Jones, Benoit Hastoy, Alejandra Tomas, Christopher A. Reissaus, Amelia K. Linnemann, Elisa D’Este, Davide Calebiro, Kai Johnsson, Tom Podewin, Johannes Broichhagen, David J. Hodson

## Abstract

The glucagon-like peptide-1 receptor (GLP1R) is a class B G protein-coupled receptor (GPCR) involved in metabolism. Presently, its visualization is limited to genetic manipulation, antibody detection or the use of probes that stimulate receptor activation. Herein, we present **LUXendin645**, a far-red fluorescent GLP1R antagonistic peptide label. **LUXendin645** produces intense and specific membrane labeling throughout live and fixed tissue. GLP1R signaling can additionally be evoked when the receptor is allosterically modulated in the presence of **LUXendin645**. Using **LUXendin645** and STED-compatible **LUXendin651**, we describe islet GLP1R expression patterns, reveal higher-order GLP1R organization including the existence of membrane nanodomains, and track single receptor subpopulations. We furthermore show that different fluorophores can confer agonistic behavior on the **LUXendin** backbone, with implications for the design of stabilized incretin-mimetics. Thus, our labeling probes possess divergent activation modes, allow visualization of endogenous GLP1R, and provide new insight into class B GPCR distribution and dynamics.

## INTRODUCTION

The glucagon-like peptide-1 receptor (GLP1R) is a secretin family class B G protein-coupled receptor (GPCR) characterized by hormone regulation.^1^ Due to its involvement in glucose homeostasis, the GLP1R has become a blockbuster target for the treatment of type 2 diabetes mellitus.^2^ The endogenous ligand, glucagon-like peptide-1 (GLP-1) is released from enteroendocrine L-cells in the gut in response to food intake,^3^ from where it travels to the pancreas before binding to its cognate receptor expressed in β-cells. Following activation, the GLP1R engages a cascade of signaling pathways including Ca^2+^, cAMP, ERK and β-arrestin, which ultimately converge on β-cell survival and the glucose-dependent amplification of insulin release.^4,5^ GLP1R is also expressed in the brain^6^ and muscle^7^ where it further contributes to metabolism *via* effects on food intake, energy expenditure, locomotion and insulin sensitivity. Despite this, GLP1R localization remains a challenge and is impeding functional characterization of GLP-1 and drug action.

Chemical biology and recombinant genetics have made available a diverse range of methods for the visualization of biological entities. Thus, classical fluorescent protein-fusions,^8^ self-labeling suicide enzymes (SNAP-, CLIP-, and Halo-tag),^9–11^ “click chemistry”^12,13^ and fluorogenic probes^14–16^ have provided unprecedented insight into the localization and distribution of their respective targets in living cells. In particular, current approaches for visualizing the GLP1R have so far relied on monoclonal antibodies (mAbs) directed against GLP1R epitopes,^17,18^ or fluorescent analogues of Exendin4(1-39),^19–21^ a stabilized form of GLP-1 and the basis for the incretin-mimetic class of drugs. Moreover, floxed mouse models exist in which Cre recombinase is driven by the *Glp1r* promoter, allowing labeling of GLP1R-expressing cells when crossed with reporter mice.^6,7^ Such methods have a number of shortcomings. Antibodies possess variable specificity^18^ and tissue penetration, and GLP1R epitopes might be hidden or preferentially affected by fixation in different cell types. Even more, fluorescent analogues of Exendin4(1-39) activate and internalize the receptor, which could confound results in live cells, particularly when used as a tool to sort purified populations (*i.e. β*-cells) for transcriptomic analysis.^22,23^ On the other hand, reporter mouse strategies possess high fidelity, but cannot account for post-translational processing, protein stability and trafficking of native receptor.^24^ Lastly, none of the aforementioned approaches are amenable to super-resolution imaging of GLP1R.

Given the wider reported roles of GLP-1 signaling in the heart,^25^ liver,^26^ immune system^2^ and brain,^27^ it is clear that new tools are urgently required to help identify GLP-1 target sites, with repercussions for drug treatment and its side effects. In the present study, we therefore set out to generate a specific probe for endogenous GLP1R detection in its native, surface-exposed state in live and fixed tissue, without receptor activation. Herein, we report **LUXendin645** and **LUXendin651**, Cy5- and SiR-conjugated far-red fluorescent antagonists with unprecedented specificity, live tissue penetration and super-resolution capability. Using our tools, we provide an updated view of GLP1R expression patterns in the islet of Langerhans, show that endogenous GLP1Rs form nanodomains at the membrane and reveal receptor subpopulations with distinct diffusion modes. Lastly, we find that installation of a TMR fluorophore unexpectedly confers potent agonist properties. As such, the **LUXendins** provide the first nanoscopic characterization of a class B GPCR, with wider flexibility for detection and interrogation of GLP1R in the tissue setting.

## RESULTS

### Design and synthesis of LUXendin555, LUXendin651 and LUXendin645

Ideally, a fluorescent probe to specifically visualize a biomolecule should have the following characteristics: straightforward synthesis and easy accessibility, high solubility, relative small size, high specificity and affinity, and a fluorescent moiety that exhibits photostability, brightness, (far-)red fluorescence with an additional two-photon cross-section. Moreover, the probe should be devoid of biological effects when applied to live cells and show good or no cell permeability, depending on its target localization. While some of these points were addressed in the past (*vide infra*), we set out to achieve this high bar by designing a highly specific fluorescent GLP1R antagonist using TMR, Cy5 and SiR fluorophores. As no small molecule antagonists for the GLP1R are known, we turned to Exendin4(9-39), a potent antagonistic scaffold amenable to modification (Fig. 1).^28^ We used solid-phase peptide synthesis (SPPS) to generate an S39C mutant,^29^ which provides a *C*-terminal thiol handle for late-stage installation of different fluorophores. As such, TMR-, Cy5- and SiR-conjugated versions were obtained by means of cysteine-maleimide chemistry, termed **LUXendin555**, **LUXendin645**, and **LUXendin651**, respectively (spectral properties are shown in Table 1, HPLC traces and HRMS characterization can be found in the SI) (Fig. 1).

**Figure 1:**
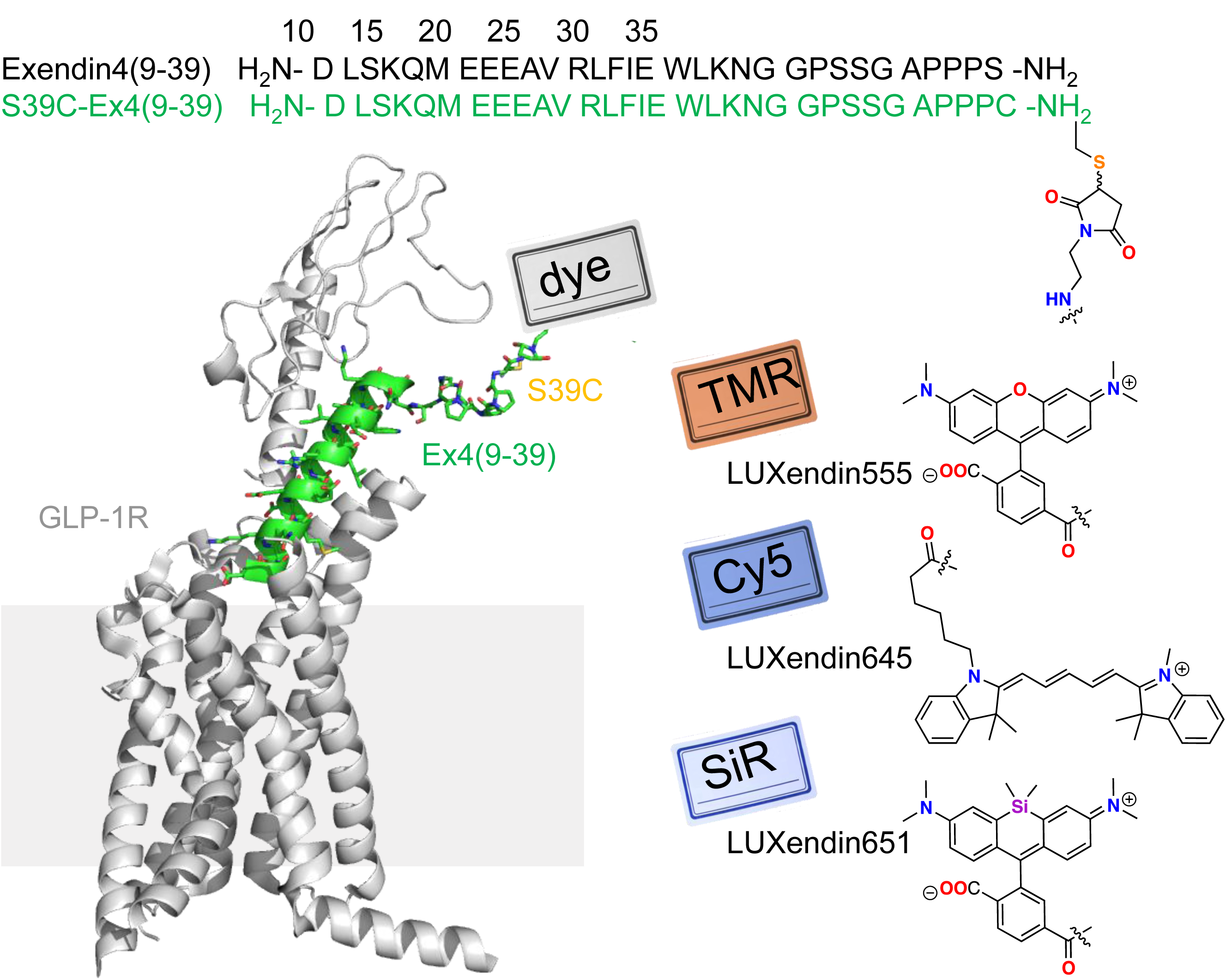
Sequence and structure of LUXendin555, LUXendin651 and LUXendin645 bound to GLP1R. **LUXendins** are based on the antagonist Exendin4(9-39), shown in complex with GLP1R. The label can be any dye, such as TMR (top), SiR (middle) or Cy5 (bottom) to give **LUXendin555, LUXendin651 and LUXendin645**, respectively. The model was obtained by using the cryo-EM structure of the activated form of GLP1R in complex with a G protein (pdb: 5VAI)^56^, with the G protein and the 8 *N*-terminal amino acids of the ligand removed from the structure while mutating S39C and adding the respective linker. Models were obtained as representative cartoons by the in-built building capability of PyMOL (Palo Alto, CA, USA) without energy optimization. Succinimide stereochemistry is unknown and neglected for clarity.

**Table 1:**
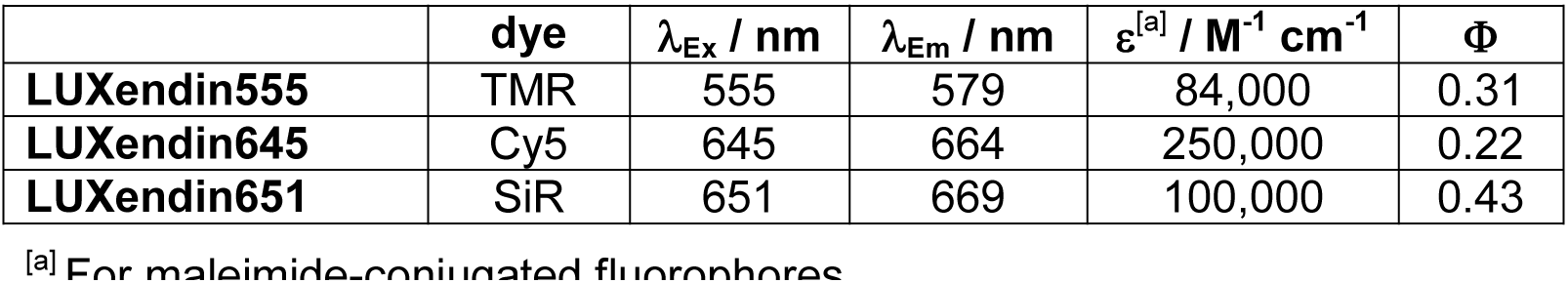
Spectral properties of GLP1R labeling probes. Maximal excitation and emission wavelengths, and quantum yields were acquired using probes dissolved at 10 µM in PBS, pH 7.4 at 21 °C.

| | dye | $\lambda_{Ex}$ / nm | $\lambda_{Em}$ / nm | $\epsilon^{[a]}$ / M<sup>-1</sup> cm<sup>-1</sup> | $\Phi$ |
| --- | --- | --- | --- | --- | --- |
| **LUXendin555** | TMR | 555 | 579 | 84,000 | 0.31 |
| **LUXendin645** | Cy5 | 645 | 664 | 250,000 | 0.22 |
| **LUXendin651** | SiR | 651 | 669 | 100,000 | 0.43 |

<sup>[a]</sup> For maleimide-conjugated fluorophores

### LUXendin645 intensely labels GLP1R in cells and tissue

GLP-1-induced cAMP production (*EC_50_*(cAMP) = 2.8 nM, 95% CI [1.5-5.2]) was similarly blocked by Exendin4(9-39) (*EC_50_*(cAMP) = 38.4 nM, 95% CI [19.0-77.8]) and its S39C mutant (*EC_50_*(cAMP) = 34.8 nM, 95% CI [18.8-64.4]) (Fig. 2a). Installation of Cy5 to produce **LUXendin645** did not affect these antagonist properties (*EC_50_*(cAMP) = 73.1 nM, 95% CI [54.9-97.5]) (Fig. 2a). As expected, addition of the GLP1R positive allosteric modulator (PAM) BETP^30^ conferred agonist activity on **LUXendin645** (*EC_50_*(cAMP) = 9.3 nM, 95% CI [2.2-40.0]), with a potency similar to Exendin4(1-39) (*EC_50_*(cAMP) = 18.3 nM, 95% CI [8.0-42.1]) (Fig. 2b).

**Figure 2:**
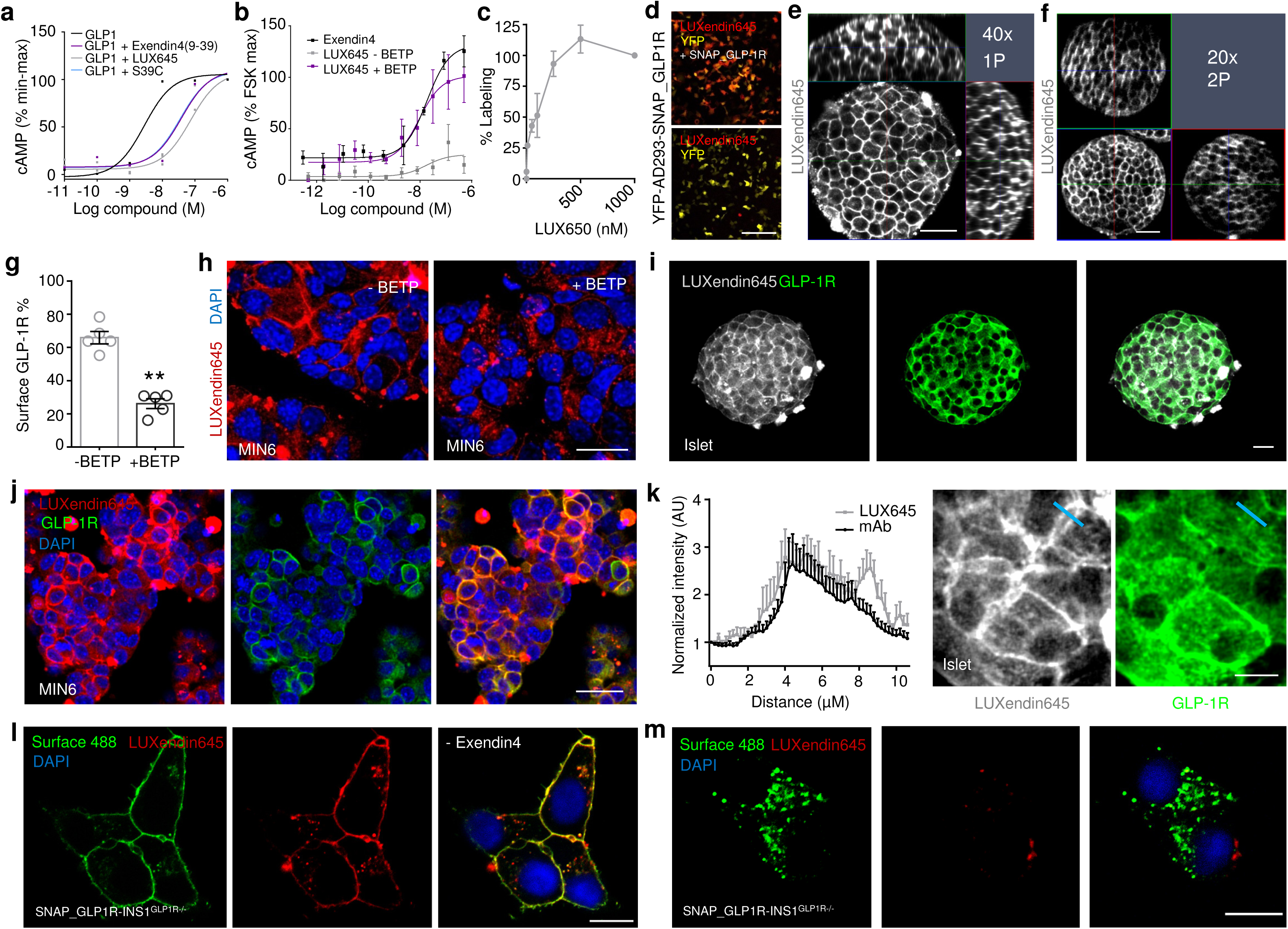
LUXendin645 binding, signaling and labeling. **a**, Exendin4(9-39), its S39C mutant and **LUXendin645** display similar antagonistic properties (n = 3 replicates). **b**, **LUXendin645** does not activate the GLP1R in CHO-K1-SNAP_GLP1R cells unless the positive allosteric modulator (PAM) BETP is present (Exendin4; +ve control) (n = 3 assays). **c**, **LUXendin645** labels CHO-K1-SNAP_GLP1R cells with a maximal labeling achieved at 100 nM. **d**, **LUXendin645** signal can be detected in YFP-AD293-SNAP_GLP1R but not YFP-AD293 cells (scale bar = 212.5 µm) (n = 3 assays). **e**, Representative confocal z-stack (1 µm steps) showing penetration of **LUXendin645** deep into a live pancreatic islet (x-y, x-z and y-z projections are shown) (n = 4 islets) (scale bar = 37.5 µm). **f**, As for (**e**), but two-photon z-stack (1 µm steps) showing the entire volume of an islet labeled with **LUXendin645** (scale bar = 37.5 µm) (n = 9 islets). **g** and **h**, GLP1R is internalized in MIN6 cells when agonist activity is conferred on **LUXendin645** using the positive allosteric modulator BETP (scale bar = 21 µm) (n = 5 images, 693-722 cells; Student’s unpaired t-test) (Bar graph shows mean ± SEM). **i** and **j**, **LUXendin645** signal can be detected even following fixation and co-localizes with a specific monoclonal antibody against the GLP1R in both islets (n = 13 islets) and MIN6 β-cells (n = 6 images, 543 cells) (scale bar = 26 µm). **k**, The superior signal-to-noise-ratio of **LUXendin645** allows more membrane detail to be visualized compared to antibody (scale bar = 12.5 µm). Representative images are shown, with a blue bar indicating the location of intensity-over-distance measures (the islet was co-stained with **LUXendin645** + antibody to allow direct comparison) (n = 13 islets). **l** and **m**, **LUXendin645** co-localizes with the SNAP label, Surface 488, in SNAP_hGLP1R-INS1^GLP1−/−^, which are deleted for the endogenous *Glp1r* (**l**). Pre-treatment with Exendin4(9-39) to internalize the GLP1R reduces **LUXendin645–**labeling (**m**) (a wash-step was used prior to application of the label) (scale bar = 10 µm) (n = 4-5 images; 57-64 cells). Mean ± SE are shown. **P<0.01.

As a first assessment of GLP1R labeling efficiency, we probed YFP-AD293-SNAP_GLP1R cells with increasing concentrations of **LUXendin645**. Maximum labeling occurred at 100 nM (Fig. 2c), in good agreement with the previously published *K_d_* = 15.8 nM of native Exendin4(9-39)^31^. **LUXendin645** was unable to label YFP-AD293 cells in which the GLP1R was absent (Fig. 2d).

We next examined whether **LUXendin645** would allow labeling of endogenous GLP1R. Following 60 min application of 50 nM **LUXendin645**, isolated islets demonstrated intense and clean labeling, which was restricted to the membrane (Fig. 2e). Using conventional confocal microscopy, we were able to detect bright staining even 60 µm into the islet (Fig. 2e). Given these results, we next attempted to penetrate deeper into the islet by taking advantage of the superior axial resolution of two-photon excitation (Fig. 2f). Remarkably, this imaging modality revealed **LUXendin645** labeling at high resolution throughout the entire volume of the islet (170 µm in this case) (Fig. 2f). Consistent with the cAMP assays, profound GLP1R internalization was detected following co-application of **LUXendin645** and BETP to MIN6 β-cells, which endogenously express the receptor (Fig. 2g, h).

### LUXendin645 allows multiplexed GLP1R detection

Demonstrating flexibility, **LUXendin645** labeling was still present following formaldehyde fixation (Fig. 2i, j). Immunostaining using a specific primary monoclonal antibody against the GLP1R revealed strong co-localization with **LUXendin645** in both islets (Fig. 2i) and MIN6 cells (Fig. 2j). Notably, **LUXendin645** displayed superior signal-to-noise-ratio and membrane resolution compared to the antibody (Fig. 2k), expected to be even better in live tissue where auto-fluorescence is less. Likewise, **LUXendin645** co-localized with SNAP-Surface 488 in SNAP_GLP1R-INS1 rat β-cells generated on an endogenous null background (Fig. 2l). Suggesting that **LUXendin645** requires the presence of surface GLP1R, labeling was markedly reduced following prior internalization with Exendin4(1-39) (Fig. 2l, m).

### LUXendin645 specifically binds the GLP1R

To further validate the specificity of **LUXendin645** labeling in primary tissue, we generated *Glp1r* knock-out mice. This was achieved using CRISPR-Cas9 genome editing to introduce a deletion into exon 1 of the *Glp1r*. The consequent frameshift was associated with absence of translation and therefore a global GLP1R knockout, termed *Glp1r*^(GE)−/−^, in which all intronic regions, and thus regulatory elements, are preserved (Fig. 3a, b). Wild-type (*Glp1r^+/+^*), heterozygous and homozygous littermates were phenotypically normal and possessed similar body weights (Fig. 3c).

**Figure 3:**
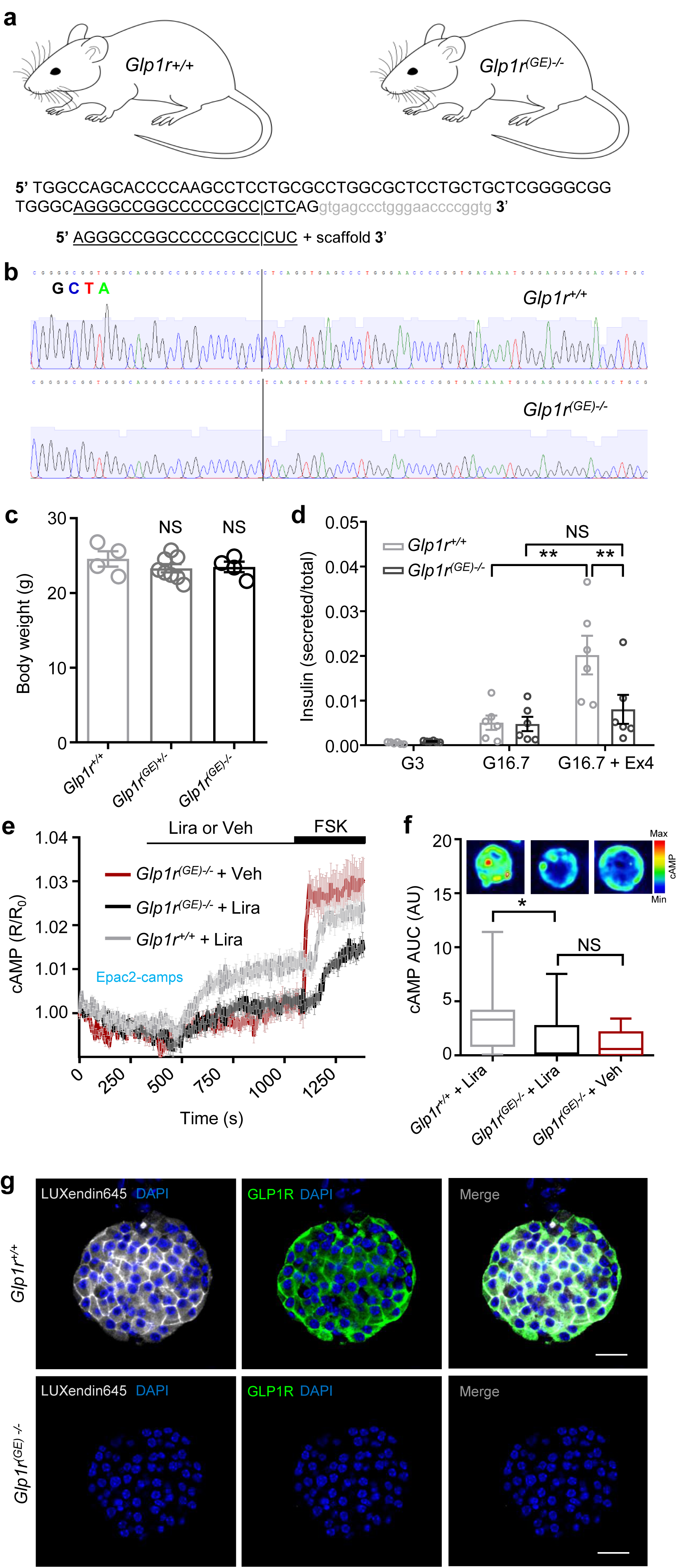
LUXendin645 is highly specific for the GLP1R. **a**, Schematic showing sgRNA targeting strategy for the production of *Glp1r^(GE)−/−^* mice. The sgRNA used targeted *Glp1r* and the double-strand break mediated by Cas9 lies within exon1 (capital letters); intron shown in gray. **b**, *Glp1r^(GE)−/−^* animals harbor a single-nucleotide deletion, as shown by sequencing traces. **c**, Body weights were similar in male 8 weeks old *Glp1r^+/+^, Glp1r^(GE)+/−^* and *Glp1r^(GE)−/−^* littermates (n = 4–8 animals; one-way ANOVA with Bonferroni’s post hoc test; F = 0.7982, DF = 2) (Bar graph shows mean ± SEM) **d**, The incretin-mimetic Exendin4(1-39) (10 nM) is unable to significantly potentiate glucose-stimulated insulin secretion in *Glp1r^(GE)−/−^* islets (n = 6 replicates; two-way ANOVA with Sidak’s post hoc test; F = 14.96, DF = 2 for *Glp1r^+/+^*, F = 2.968, DF = 2 for *Glp1r^(GE)−/−^*) (Bar graph shows mean ± SEM) **e**, Liraglutide (Lira) does not stimulate cAMP beyond vehicle (Veh) control in *Glp1r^(GE)−/−^* islets, measured using the FRET probe Epac2-camps (traces represent mean ± SEM) (n = 14–17 islets). **f**, cAMP area-under-the-curve (AUC) quantification showing absence of significant Liraglutide-stimulation in *Glp1r^(GE)−/−^* islets (n = 14–17 islets; Kruskal-Wallis test with Dunn’s post hoc test; Kruskal-Wallis statistic = 7.6, DF = 2) (Box and Whiskers plot shows min-max and median) (representative images displayed above each bar). **g**, **LUXendin645** and GLP1R antibody labeling is not detectable in *Glp1r^(GE)−/−^* islets (scale bar = 40 µm) (n = 12–14 islets for each genotype). *P<0.05, **P<0.01 and NS, non-significant.

Confirming successful GLP1R knock-out, insulin secretion assays in islets isolated from *Glp1r^(GE)−/−^* mice showed intact responses to glucose, but absence of Exendin4(1-39)-stimulated insulin secretion (Fig. 3d). Reflecting this finding, the incretin-mimetic Liraglutide was only able to stimulate cAMP rises in islets from wild-type (*Glp1r^+/+^*) littermates, measured using the FRET probe Epac2-camps (Fig. 3e, f). As expected, immunostaining with monoclonal antibody showed complete absence of GLP1R protein (Fig. 3g). Suggesting that **LUXendin645** specifically targets GLP1R, with little to no cross-talk from glucagon-receptors,^32^ signal could not be detected in *Glp1r^(GE)−/−^* islets (Fig. 3g).

Together, these data provide strong evidence for a specific mode of **LUXendin645** action *via* the GLP1R.

### LUXendin645 highlights weak GLP1R expression

Previous approaches have shown low abundance of *Glp1r* transcripts in the other major islet endocrine cell type, *i.e.* glucagon-secreting α-cells.^7,33^ This is associated with detection of GLP1R protein in ∼1-10% of cells,^7,34^ providing an excellent testbed for **LUXendin645** sensitivity. Studies in intact islets showed that **LUXendin645** labeling was widespread in the islet and well co-localized with insulin immunostaining (Fig. 4a). However, **LUXendin645** could also be seen on membranes very closely associated with α-cells and somatostatin-secreting δ-cells (Fig. 4b, c), similarly to results obtained with GLP1R mAb. Due to the close apposition of β-, α- and δ-cell membranes, we were unable to accurately assign cell-type specificity to **LUXendin645**. Instead, using cell clusters plated onto coverslips, we could better discern **LUXendin645** labeling, revealing GLP1R expression in 18 ± 6% of α-cells (Fig. 4d–f), higher than that shown before using antibodies^19,34^ and reporter genes^7^. Notably, GLP1R-expressing α-cells tended to adjoin, whereas those without the receptor were next to β-cells. Confirming previous findings, a majority (86 ± 3%) of β-cells were positive for **LUXendin645** (Fig 4d-f).^7,19^

**Figure 4:**
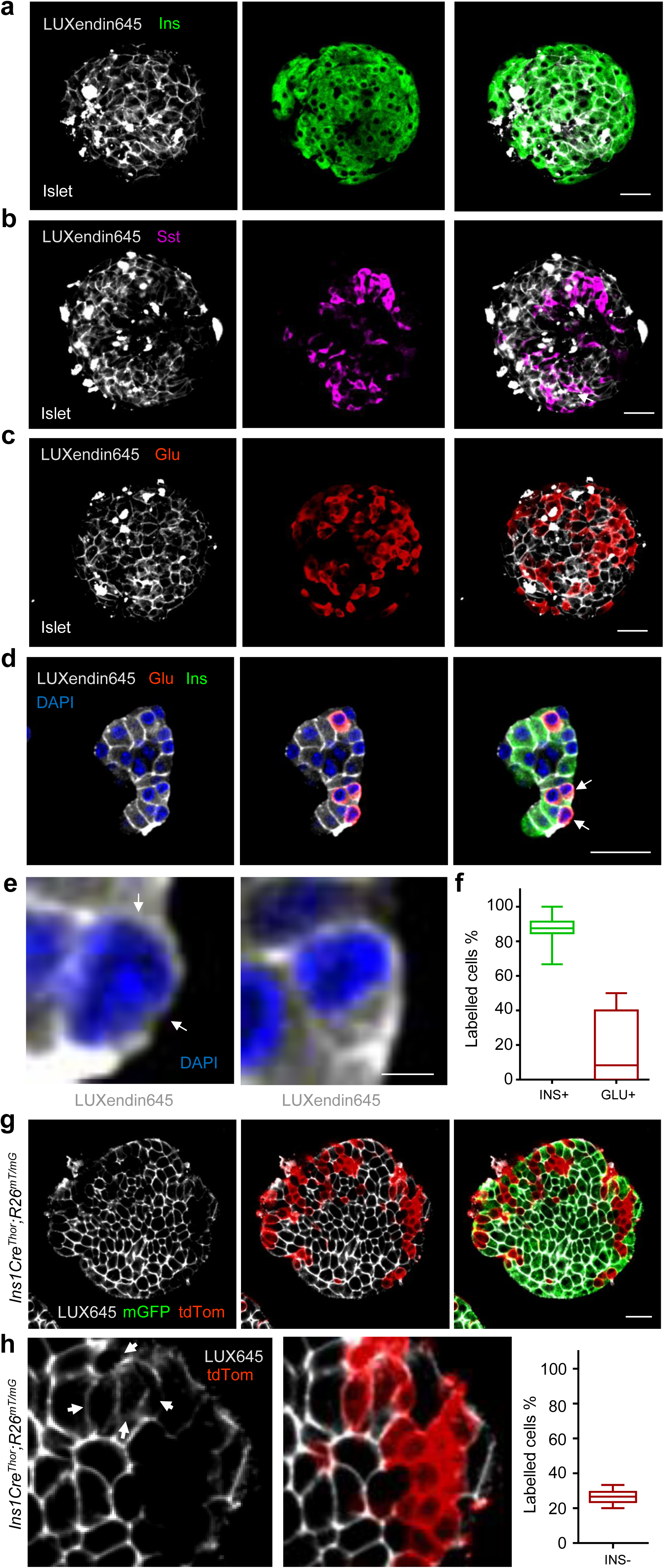
LUXendin645 reveals GLP1R expression in a subpopulation of α-cells. **a-c**, **LUXendin645** labeling is widespread throughout the intact islet, co-localizing predominantly with β-cells (**a**) and δ-cells (**b**), but less so with α-cells (**c**) stained for insulin, somatostatin and glucagon, respectively (n = 7–9 islets) (scale bar = 26 µm). **d**, Following dissociation of islets into cell clusters, **LUXendin645** labeling can be more accurately quantified (arrows highlight cells selected for zoom-in) (scale bar = 26 µm). **e**, Zoom-in of (**d**) showing a **LUXendin645**-(left) and **LUXendin645**+ (right) α-cell (arrows highlight non-labeled cell membrane, which is not bounded by a β-cell) (scale bar = 26 µm). **f**, Box-and-whiskers plot showing proportion of β-cells (INS) and α-cells (GLU) co-localized with **LUXendin645** (n = 5–6 images, 12 cell clusters) (Max-min shown together with the median). **g**, *Ins1Cre^Thor^;R26^mT/mG^* dual fluorophore reporter islets express tdTomato until Cre-mediated replacement with mGFP, allowing identification of β-cells (∼80% of the islet population) and non-β-cells for live imaging (scale bar = 26 µm). **LUXendin645** highlights GLP1R expression in nearly all β-cells but relatively few non-β-cells (n = 24 islets, 809 cells). **h**, As for (**g**), but a zoom-in showing GLP1R expression in some non-β-cells (left) together with quantification (right) (arrows show **LUXendin645-**labeled non-β cells) (scale bar = 5 µm) (Box and Whiskers plot shows min-max and median).

We wondered whether fixation required for immunostaining might increase background fluorescence such that GLP1R detection specificity was reduced. To circumvent this, studies were repeated in live islets where **LUXendin645** signal was found to be much brighter and background almost non-existent. GLP1R was detected in 26.2 ± 1.1 % of non-β-cells (Fig 4g, h) using *Ins1Cre^Thor^;R26^mTmG^* reporter mice in which β-cells are labeled green and all other cell types are labeled red following Cre-mediated recombination. Once adjusted for the previously reported GLP1R expression in δ-cells (assuming 100%), which constitute ∼20% of the insulin-negative islet population,^35^ this leaves ∼6% of GLP1R+ α-cells. This was not an artefact of optical section, since two-photon islet reconstructions showed similar absence of **LUXendin645** staining in discrete regions near the surface (where α-cells predominate) (Movie S1).

### LUXendin645 and Luxendin651 reveal higher-order GLP1R organization

By combining **LUXendin645** with Super-Resolution Radial Fluctuations (SRRF) analysis,^36^ GLP1R could be imaged at super-resolution using streamed images (∼ 1000) from a conventional widefield microscope (Fig. 5a). To image endogenous GLP1R at < 100 nm lateral resolution, we combined STED nanoscopy with **LUXendin651**, which bears silicon rhodamine (SiR) instead of Cy5. **LUXendin651** produced bright labeling of wild-type but not *Glp1r^(GE)^*^−/−^ islets, with an identical distribution to **LUXendin645** (Supplementary Fig. S1). Incubation of MIN6 cells with **LUXendin651** and subsequent fixation allowed STED imaging of the endogenous GLP1R with a FWHM = 70±10 nm (Fig. 5b, c). STED snapshots of MIN6 β-cells revealed GLP1R distribution with unprecedented detail: receptors were not randomly arranged but rather tended to organize into nanodomains with neighbors (Fig. 5b, c). This was confirmed using the F- and G-functions, which showed a non-random and more clustered GLP1R distribution (Fig. 5d, e). Differences in GLP1R expression level and pattern could clearly be seen between neighboring cells with a subpopulation possessing highly concentrated GLP1R clusters (Fig. 5f).

**Figure 5:**
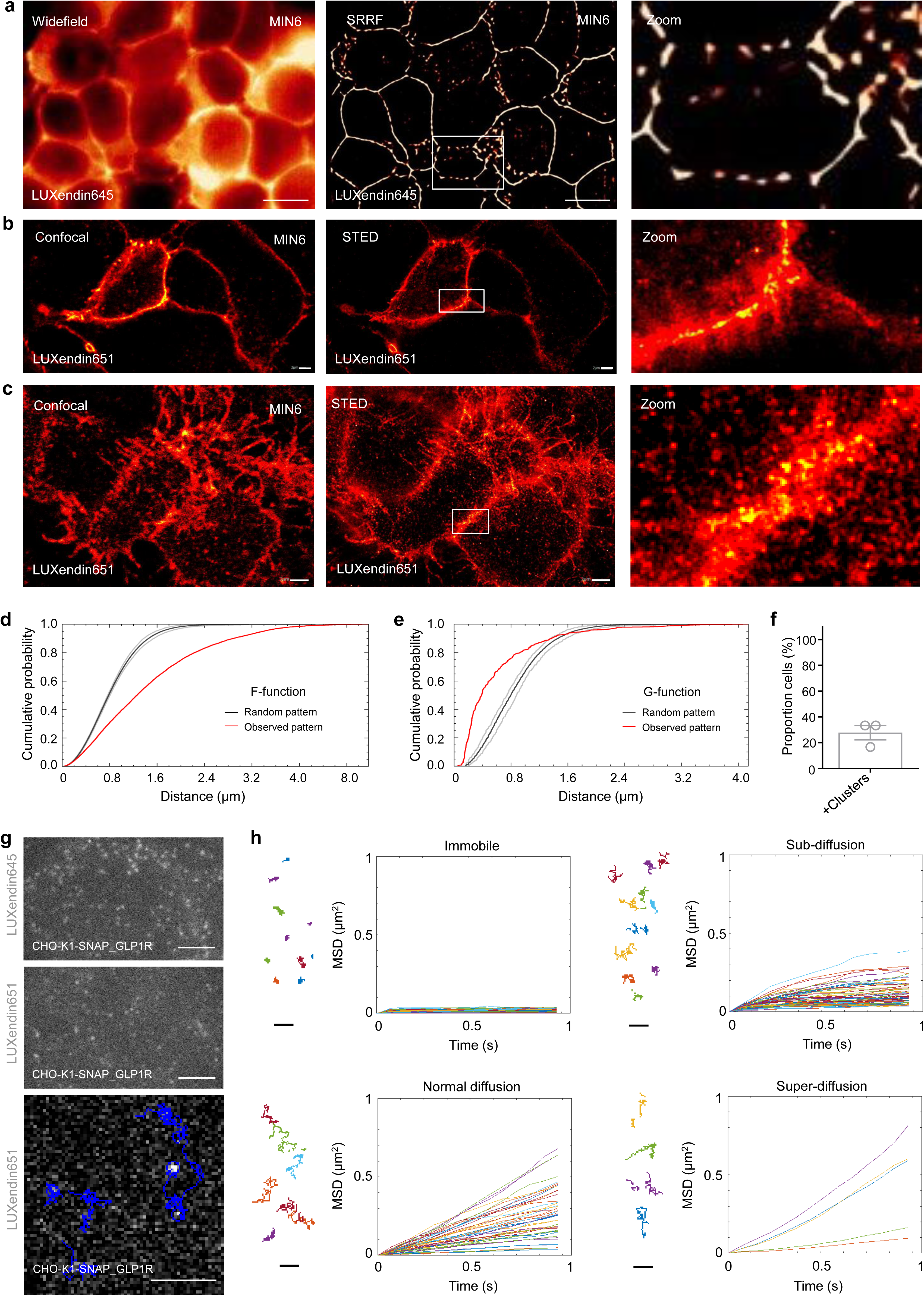
LUXendin651 and LUXendin645 allow nanoscopic detection of GLP1R distribution and dynamics. **a**, **LUXendin645** allows super-resolution snapshots of MIN6 β-cells using widefield microscopy combined with Super-Resolution Radial Fluctuations (SRRF) (n = 3 images) (scale bar = 10 µm). **b** and **c**, Confocal and STED snapshots of endogenous GLP1R in **LUXendin651**-treated MIN6 cells at ∼ 50 nm axial resolution. Note the presence of punctate GLP1R expression as well as aggregation/clustering in images captured above (**b**) and close to the coverslip (**c**) using STED microscopy (n = 3 images, 15 cells) (scale bar = 2 µm). **d** and **e**, Spatial analysis of GLP1R expression patterns using the F-function (**d**) and G-function (**e**) show a non-random distribution (red line) versus a random model (black line; 95% confidence interval shown). **f**, Approximately 1 in 4 MIN6 β-cells possess highly concentrated GLP1R clusters (Bar graph shows mean ± SEM) (n = 3 images, 15 cells). **g**, Single molecule microscopy and tracking of **LUXendin645**- and **LUXendin651**-labeled GLP1R (n = 2 movies) (scale bar = 3 µm). **h**, Mean square displacement (MSD) analysis showing different GLP1R diffusion modes (representative trajectories are displayed) (scale bar = 1 µm).

Finally, to test whether **LUXendin645** and **LUXendin651** would be capable of tracking GLP1Rs in live cells, we performed single-molecule microscopy experiments in which individual receptors labeled with either fluorescent probe were imaged by total internal reflection fluorescence (TIRF) microscopy.^37,38^ Both probes allowed GLP1R to be tracked at the single-molecule level in CHO-K1-SNAP_GLP1R cells, but brightness and bleaching precluded longer recordings with **LUXendin645** (Fig. 5g and Supplementary Movies S2, S3). By combining single-particle tracking with **LUXendin651**, we were able to show that most GLP1Rs diffuse rapidly at the membrane (Fig. 5g and Supplementary Movie S4). However, a mean square displacement (MSD) analysis^37^ revealed a high heterogeneity in the diffusion of GLP1Rs on the plasma membrane, ranging from virtually immobile receptors to some displaying features of directed motion (superdiffusion) (Fig. 5h).

### Altering fluorophore to produce LUXendin555 confers different ligand behavior

Lastly, we explored whether swapping the far-red Cy5/SiR for a TMR dye would be tolerated to obtain a spectrally orthogonal probe, termed **LUXendin555**. Labeling was detected in YFP-AD293-SNAP_GLP1R (Fig. 6a) but not YFP-AD293 cells (Fig. 6b). However, we noticed a more punctate **LUXendin555** staining pattern when viewed at high-resolutions (Fig. 6c). Further experiments with MIN6 cells and islets showed similar internalization of the GLP1R (Fig. 6d), suggesting that **LUXendin555** acts as an agonist, presumably via interactions mediated by the ectodomain. This was confirmed using cAMP assays where **LUXendin555** was found to potently activate GLP1R signaling (*EC_50_*(cAMP) = 129.8 nM; 95% CI = 56.9–296.2) (Fig. 6e). Intriguingly, **LUXendin555** potency could be further increased using a PAM (*EC_50_*(cAMP) = 28.4 nM; 95% CI = 11.3-71.8) (Fig. 6f), suggesting a unique binding conformation at the orthosteric site compared to agonists such as Exendin4(1-39), which cannot be allosterically-modulated.^30^ As for the other probes, **LUXendin555** was unable to label *Glp1r^(GE)^*^−/−^ islets (Fig. S2).

**Figure 6:**
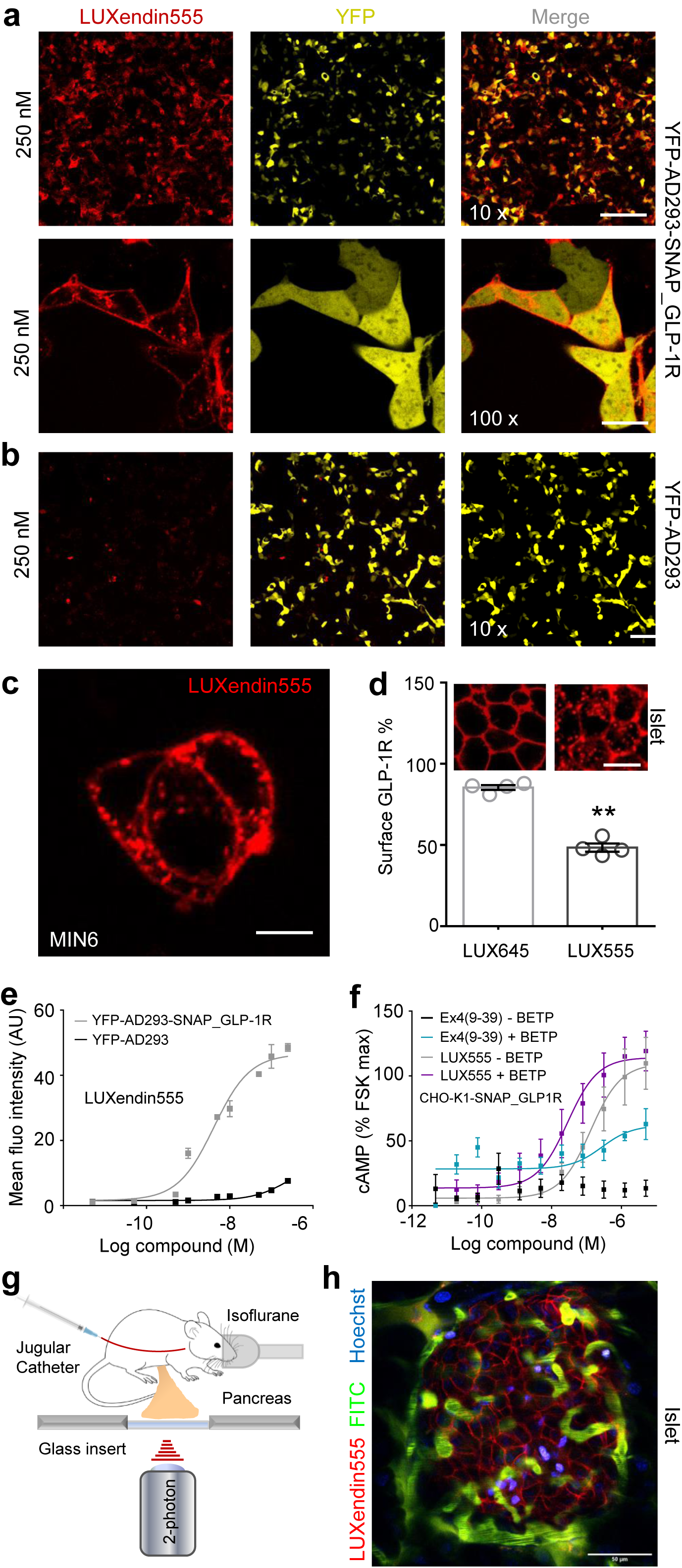
LUXendin555 displays agonist properties and allows *in vivo* labeling of islets. **a** and **b**, **LUXendin555** labels YFP-AD293_SNAP-GLP1R (**a**) but not YFP-AD293 (**b**) controls (n = 3–4 assays) (10x scale bar = 213 µm; 100x scale bar = 21 µm). **c**, High resolution snapshot of **LUXendin555**-labeling in MIN6 β-cells showing a punctate staining pattern in the cytosol (n = 8 images, 142 cells) (scale bar = 9 µm). **d**, Surface GLP1R expression is reduced in **LUXendin555**-compared to **LUXendin645**-treated islets (representative images shown above each bar) (n = 5 islets; Student’s unpaired t-test) (Bar graph shows mean ± SEM) (scale bar = 17 µm). **e**, **LUXendin555** potently increases cAMP levels in YFP-AD293_SNAP-GLP1R but not YFP-AD293 cells (n = 3 assays). **f**, Allosteric modulation with BETP increases agonist activity of **LUXendin555** (n = 3 assays). **g**, Schematic depicting the two-photon imaging set up for visualization of the intact pancreas in mice. **h**, Representative image showing that **LUXendin555** labels cell membranes in an islet surrounded by the vasculature *in vivo* (n = 2 islets from 1 mouse) (scale bar = 50 µm). Mean ± SE are shown. **P<0.01.

### LUXendins label islets *in vivo*

We thought that the high quantum yield of TMR, coupled with good two-photon cross-section and agonistic behaviour might suit **LUXendin555** well to *in vivo* imaging where maintenance of normoglycemia under anaesthesia can be an advantage for some experiments. Two-photon imaging was applied to an anaesthetized mouse to allow visualization of the intact pancreas, exposed through an abdominal incision (Fig. 6g). Vessels and nuclei were first labeled using FITC-dextran and Hoechst before injecting **LUXendin555** intravenously. Labeling occurred rapidly within 5 min post-injection, produced intense membrane staining confined to the islet where GLP1R is expressed (Fig. 6h), and normoglycemia was maintained (250 mg/dl). No obvious internalization could be seen, most likely reflecting the time of exposure to **LUXendin555**, as well as the concentration achieved *in vivo* at the islet.

## DISCUSSION

In the present study, we synthesize and validate far-red fluorescent labels, termed **LUXendins** for the real-time detection of GLP1R in live cells. Nanomolar concentrations of **LUXendin645** and **LUXendin651** led to intense membrane-labeling of the GLP1R, with best in class tissue penetration and signal-to-noise ratio, as well as super-resolution capability. Notably, **LUXendin645** and **LUXendin651** did not activate the GLP1R unless agonist activity is conferred with the widely-available PAM BETP. **LUXendin645** and **LUXendin651** are highly specific, as shown using a novel CRISPR-Cas9 mouse line lacking GLP1R expression. Lastly, the analogous compound **LUXendin555** bearing a different fluorophore unusually displays agonistic activity, expanding the color palette and activity profile without changing the peptidic pharmacophore.

Compared to present chemical biology approaches, **LUXendins** possess a number of advantages for GLP1R labeling, which generally rely on Exendin4(1-39) labeled with for instance Cy3, Cy5 or FITC.^19–21,30^ Firstly, the use of an antagonist encourages receptor recycling back to the membrane and retains receptor at the cell surface, which likely increases detection capability. Secondly, the GLP1R is not activated, meaning that results can be interpreted in the absence of potentially confounding cell signaling, such as that expected with agonists.^19^ Thirdly, Cy5 occupies the far-red range, leading to less background fluorescence, increasing depth penetration due to reduced scatter, and avoiding the use of more phototoxic wavelengths.^39^ Together, these desirable properties open up the possibility to image expression and trafficking of native GLP1R over extended periods of time, when **LUXendins** are used in conjunction with a PAM.

To test the specificity of **LUXendins**, we used CRISPR-Cas9 genome-editing to globally knock out the GLP1R in mice. Protein deletion was confirmed by absence of detectable GLP1R signal following labeling with monoclonal antibody, **LUXendin555**, **LUXendin645** and **LUXendin651**. While *Glp1r*^−/−^ animals already exist and have made important contributions to our understanding of incretin biology, they were produced using a targeted mutation to replace exons encoding transmembrane regions 1 and 3 (encoded by exons 5 and 7), presumably leading to deletion of the introns in between (∼6.25 kb).^40^ By contrast, *Glp1r*^(GE)−/−^ mice possess intact introns. Since introns contain regulatory elements, such as distant-acting enhancers^41^, miRNAs^42^ and lncRNAs,^43^ their loss in transgenic knockouts could have wider influence on the transcriptome. GLP1R knock-out mice might therefore be useful alongside conventional approaches for validating GLP1R reagents, including antibodies, agonist and antagonist, and derivatives thereof.

Demonstrating the excellent sensitivity of the Cy5-linked **LUXendin645** in particular, we were able to detect GLP1R in ∼6-18% of α-cells. Understanding α-cell GLP1R expression patterns is important because incretin-mimetics reduce glucagon secretion,^44^ which would otherwise act to aggravate blood glucose levels. However, previous studies using antibodies, reporter animals and agonist-fluorophores have shown only ∼1-10% GLP1R expression in mouse and rat α-cells, in line with the low transcript abundance^7,19,33,45^, despite reports that GLP-1 can directly suppress glucagon release.^34,46^ Our data are in general concordance with these findings, but demonstrate an increase in detection capability for native GLP1R. This improvement is likely related to the superior SNR of **LUXendin645** compared to mAb and agonist-fluorophore, increasing the ability to resolve relatively low GLP1R levels. A recent report showed GLP1R expression in ∼80% of α-cells using a novel antibody raised against the *N*-terminal region, with both membrane and cytosolic staining evident^47^. While the reasons for this discrepancy are unknown, it should be noted that **LUXendin645** binds the orthosteric site and so reports the proportion of GLP1R that is “signaling competent”.^7,19,32^.

Since **LUXendin645** showed excellent signal-to-noise ratio using conventional epifluorescence, it was highly amenable to SRRF analysis. As such, **LUXendin645** and its congeners open up the possibility to image the GLP1R at super-resolution using simple widefield microscopy available in most laboratories. For stimulated emission depletion (STED) microscopy experiments, Cy5 was replaced with SiR to give **LUXendin651**. STED imaging showed that endogenous GLP1R possess a higher structural order, namely organization into nanodomains at the cell membrane. The presence of nanodomains under non-stimulated conditions might reflect differences in palmitoylation, which has recently been shown to influence GLP1R membrane distribution in response to agonists.^48^ Notably, a subpopulation of β-cells appeared to possess highly-concentrated GLP1R clusters. It will be important in the future to investigate whether this is a cell autonomous heterogenous trait, or instead reflects biased orientation of membranes toward specific β-cells. Lastly, both **LUXendin645** and **LUXendin651** allowed GLP1Rs to be imaged in live cells by single-molecular microscopy, revealing variability in their diffusion at the plasma membrane. Particle tracking analyses segregated GLP1R into four different populations based upon diffusion mode, in keeping with data from beta adrenergic receptors.^37^ Together, these experiments provide the first super-resolution characterization of a class B GPCR and suggest a degree of complexity not readily appreciated with previous approaches.

Intriguingly, we saw that swapping Cy5 for a TMR moiety to give **LUXendin555** completely changed the pharmacological behavior. The reasons for this are unknown, but we speculate that the rhodamine scaffold engages a secondary binding site in the GLP1R ectodomain, leading to activation. This finding is remarkable because it suggests that the agonist antagonist switch that occurs following removal of eight *N*-terminal amino acids (as physiologically mediated by the protease DPP-4)^49^ can be counteracted simply by installing a *C*-terminal linked rhodamine fluorophore, with implications for the design of more stable GLP1R activators. More generally, this shift in compound behaviour following a fluorophore modification serves as another instructive example for the thorough validation of all new chemical labels.^50^ Nonetheless, **LUXendin555** possessed advantageous properties for *in vivo* imaging including maintenance of relatively stable glycemia, good two-photon cross-section and high quantum yield.

In summary, we provide a comprehensively-tested and unique GLP1R detection toolbox consisting of far-red antagonist labels, **LUXendin645** and **LUXendin651**, an agonist **LUXendin555**, and knockout *Glp1r^(GE)−/−^* animals. Using these freely-available probes, we provide an updated view of GLP1R organization, with relevance for the treatment of complex metabolic diseases such as obesity and diabetes, as well as production of more stable GLP1R activators. Thus, the stage is set for visualizing GLP1R in various tissues using a range of imaging techniques, as well as the production of novel peptidic labels and agonists.

## METHODS

### Synthesis

Solid-phase peptide synthesis of S39C-Exendin4(9-39) was performed as previously reported.^29^ Maleimide-conjugated-6-TMR, −6-SiR and −Cy5 were obtained by TSTU activation of the corresponding acids and reaction with 1-(2-amino-ethyl)-pyrrole-2,5-dione (TFA salt, Aldrich). Fluorophore coupling *via* thiol-maleimide chemistry to peptides was performed in PBS. All compounds were characterized by HRMS and purity was assessed to be >95% by HPLC. Extinction coefficients were based upon known manufacturer bulk material measures for TMR-Mal, Cy5-Mal (both Lumiprobe) and SiR-Mal (Spirochrome). Details for synthesis including further characterization of all **LUXendins** are detailed in the Supporting Information. **LUXendin555, LUXendin651** and **LUXendin645** are freely available for academic use upon request.

### Cell culture

AD293 cells (Agilent) were maintained in Dulbecco’s Modified Eagles medium (DMEM) supplemented with 10% fetal calf serum (FCS), 1% *L*-glutamine and 1% penicillin/streptomycin. CHO-K1 cells (a kind gift from Dr Ben Jones, Imperial College London) stably expressing the human SNAP_GLP1R (Cisbio) (CHO-K1-SNAP_GLP1R) were maintained in DMEM supplemented with 10% FCS, 1% penicillin/streptomycin, 500 μg/mL G418, 5 mM *D*-glucose, 10 mM HEPES and 1% nonessential amino acids. MIN6 β-cells (a kind gift from Prof. Jun-ichi Miyazaki, Osaka University) were maintained in DMEM supplemented with 15% FCS, 25 mM *D*-glucose, 71 μM BME, 2 mM *L*-glutamine, 100 U/mL penicillin, and 100 μg/mL streptomycin. INS1 832/3 CRISPR-deleted for the endogenous GLP1R locus (a kind gift from Dr. Jacqui Naylor, MedImmune)^51^ were transfected with human SNAP_GLP1R, before FACS of the SNAP-Surface488-positive population and selection using G418.^48^ The resulting SNAP_GLP1R_INS1^GLP1R−/−^ cells were maintained in RPMI-1640 supplemented with 10% FBS, 10 mM HEPES, 2 mM *L*-glutamine, 1 mM pyruvate, 72 µM β-mercaptoethanol, 1% penicillin/streptomycin and 500 μg/mL G418.

### Animals

*Glp1r*^(GE)−/−^: CRISPR-Cas9 genome-editing was used to introduce a single base pair deletion into exon 1 of the *Glp1r* locus. Fertilized eggs of female Cas9-overexpressing mice (strain *Gt(ROSA)26Sor^tm1.1(CAG-cas9*,–EGFP)Fezh^*/J) were harvested following super-ovulation. Modified single-guide RNA (Synthego) targeting exon 1 of *Glp1r* and a single-stranded repair-template were injected at 20 ng/µl into the pronucleus of embryos at the 1-cell stage. In culture, 80% of embryos reached the 2-cell stage and were transplanted into surrogate mice. The targeted locus of offspring was analyzed by PCR and sequencing. Besides the insertion of the repair template, deletions of up to 27 nucleotides could be detected in 2 out of 6 offspring. Design of the repair template will be described elsewhere. Off-target sites were predicted using the CRISPR Guide Design Tool (crispr.mit.edu). Loci of the top ten off-target hits were amplified by PCR and analyzed *via* Sanger sequencing. Founder animals carrying alleles with small deletions were backcrossed to wild type animals (strain C57BL/6J) for 1–2 generations to outbreed affected off-targets and then bred to homozygosity. Animals with the larger deletion of 27 nucleotides were not taken forward, as GLP1R protein was still present. Animals were born in Mendelian ratios and genotyping was performed using Sanger sequencing. Animals were bred as heterozygous pairs to ensure *Glp1r*^+/+^ littermates. *Glp1r*^(GE)−/−^ animals are freely available for academic use, subject to a Material Transfer Agreement.

*Ins1Cre^Thor^;R26^mT/mG^:* To allow identification of β- and non-β-cells, *Ins1Cre^Thor^ a*nimals with Cre knocked-in at the *Ins1* locus (strain B6(Cg)-*Ins1^tm1.1(cre)Thor^*/J) were crossed with *R26^mT/mG^* reporter mice (strain B6.129(Cg)-*Gt(ROSA)26Sor^tm4(ACTB-tdTomato,-EGFP)Luo^*/J). Cre-dependent excision of the floxed allele results in deletion of tdTomato, expression of membrane-localized GFP and thus identification of recombined and non-recombined cells.

All studies were performed with 6-12 week old male and female animals, and regulated by the Animals (Scientific Procedures) Act 1986 of the U.K. Approval was granted by the University of Birmingham’s Animal Welfare and Ethical Review Body.

### Islet isolation

Islets were isolated from male and female *Glp1r^(GE)−/−^* and *Ins1Cre^Thor^;R26^mT/mG^* mice, as well as CD1 wild-type animals, maintained under specific-pathogen free conditions, with *ad lib* access to food and water. Briefly, animals were humanely euthanized before injection of collagenase 1 mg/mL (Serva NB8) into the bile duct. Following removal of the inflated pancreas and digestion for 12 min at 37 °C, islets were separated using a Histopaque (Sigma-Aldrich) gradient. Islets were cultured in RPMI medium containing 10% FCS, 100 units/mL penicillin, and 100 μg/mL streptomycin.

### Binding and potency assays

Binding assays were performed in transiently-transfected YFP-AD293-SNAP_GLP1R cells (using PolyJet reagent; SignaGen). Increasing concentrations of compound were applied for 60 min, before imaging using a Zeiss LSM880 meta-confocal microscope configured with GaAsP detectors and 10×/0.45 W, 40x/1.00 W and 63×/1.20 W objectives. YFP, TMR (**LUXendin555**) and Cy5 (**LUXendin645**) were excited using λ = 514 nm, λ = 561 nm and λ = 633 nm lasers, respectively. Emitted signals were captured at λ = 519–574 nm, λ = 570– 641 nm and λ = 638-759 nm for YFP, TMR (**LUXendin555**) and Cy5 (**LUXendin645**), respectively. Control experiments were performed in YFP-AD293-SNAP cells, as above.

Potency for cAMP generation and inhibition was tested in heterologous expression systems, comprising either stable CHO-K1-SNAP_GLP1R cells or transiently-transfected YFP-AD293-SNAP_GLP1R cells, as previously described.^29^ Briefly, cells were incubated with increasing concentrations of compound +/- allosteric modulator for 30 min, before harvesting, lysis and measurement of cAMP using cAMP-Glo^TM^ Assay (Promega), according to the manufacturer’s instructions. *EC_50_* values were calculated using log concentration-response curves fitted with a three-parameter equation.

### Live imaging

Islets were incubated for 1 h at 37 °C in culture medium supplemented with either 100-250 nM **LUXendin555**, 50-100 nM **LUXendin645** or 100 nM **LUXendin651**, based upon binding assays. Islets were imaged using either a Zeiss LSM780 or LSM880 microscope, as above (**LUXendin651** was imaged as for **LUXendin645**). *Ins1Cre^Thor^;R26^mT/mG^* islets were excited at λ = 488 nm (emission, λ = 493-555 nm) and λ = 561 nm (emission, λ = 570-624 nm) for mGFP and tdTomato, respectively. Two-photon imaging of **LUXendin645** was performed using a Zeiss LSM 880 NLO equipped with a Spectra-Physics Insight X3 femtosecond-pulsed laser and 20x/1.00 W objective. Excitation was performed at λ = 800 nm and emitted signals detected at λ = 638-759 nm.

### cAMP imaging

Islets were transduced with adenovirus harboring the FRET sensor, Epac2-camps, before imaging using a Crest X-Light spinning disk system coupled to a Nikon Ti-E base and 10x/0.4 NA objective. Excitation was delivered at λ = 430–450 nm using a Lumencor Spectra X light engine. Emitted signals were detected at λ = 460–500 and λ = 520–550 nm for Cerulean and Citrine, respectively, using a Photometrics Delta Evolve EM-CCD. Imaging was performed in HEPES-bicarbonate buffer, containing (in mmol/L) 120 NaCl, 4.8 KCl, 24 NaHCO_3_, 0.5 Na_2_HPO_4_, 5 HEPES, 2.5 CaCl_2_, 1.2 MgCl_2_, and 3–17 *D*-glucose. Vehicle (H_2_0), Exendin4(1-39) (10-20 nM) or Liraglutide (10 nM) were applied at the indicated time points, with forskolin (10 µM) acting as a positive control.

### Immunostaining

**LUXendin555**- or **LUXendin645**-treated cells or tissue were fixed for 60 min in 4% paraformaldehyde. Primary antibodies were applied overnight at 4 °C in PBS + 0.1% Triton

+ 1% BSA. Secondary antibodies were applied in the same buffer for 1 h at room temperature, before mounting on slides using Vectashield Hardset containing DAPI. Primary antibodies were mouse monoclonal anti-GLP1R 1:30 (Iowa DHSB; mAb #7F38), rabbit anti-insulin 1:500 (Cell Signaling Technology, #3014), mouse monoclonal anti-glucagon 1:2000 (Sigma-Aldrich, #G2654) and mouse anti-somatostatin 1:5000 (Invitrogen, #14-9751-80). Secondary antibodies were goat anti-mouse Alexa Fluor 568 and donkey anti-rabbit DyLight 488 1:1000. Images were captured using an LSM880 meta-confocal microscope. Alexa Fluor 488 and Alexa Fluor 568 were excited at λ = 488 nm and λ = 568 nm, respectively. Emitted signals were detected at λ = 500–550 nm (Alexa Fluor 488) and λ = 519–574 nm (Alexa Fluor 568).

### Super-resolution microscopy

*SRRF:* MIN6 were treated with **LUXendin645** before fixation and mounting on slides using Vectashield Hardset containing DAPI. Imaging was performed using a Crest X-Light spinning disk system in bypass (widefield) mode. Excitation was delivered at λ = 640/30 nm through a 63x/1.4 NA objective using a Lumencor Spectra X light engine. Emission was collected at λ = 700/75 nm using a Photometrics Delta Evolve EM-CDD. A 1000 image sequence was captured (∼ 2 min) before offline super resolution radial fluctuation (SRRF) analysis to generate a single super-resolution snapshot using the NanoJ plugin for ImageJ (NIH).^36^ *Stimulated emission depletion (STED) microscopy*: MIN6 cells were treated with 100, 200 and 400 nM **LUXendin651** before fixation (4% paraformaldehyde, 20 min). Cells were mounted in Mowiol supplemented with DABCO and imaged on an Abberior STED 775/595/RESOLFT QUAD scanning microscope (Abberior Instruments GmbH, Germany) equipped with STED lines at λ = 595 and λ = 775 nm, excitation lines at λ = 355 nm, 405 nm, 485 nm, 561 nm, and 640 nm, spectral detection, and a UPlanSApo 100x/1.4 oil immersion objective lens. Following excitation at λ = 640 nm, fluorescence was acquired in the spectral window λ = 650-800 nm. Deconvolution was performed with Richardson-Lucy algorithm on Imspector software. FWHM was measured on raw data and calculated using OriginPro 2017 software with Gaussian fitting (n=15 profiles). Spatial GLP1R expression patterns were analyzed using the F- and G-functions, where F = distance between an object of interest and its nearest neighbor, and G = distance from a given position to the nearest object of interest (FIJI Spatial Statistic 2D/3D plugin).^52^ Both measures were compared to a random distribution of the same measured objects, with a shift away from the mean +/- 95% confidence intervals indicating a non-random or clustered organization (*i.e.* more space or smaller distance between objects).

*Single-molecule microscopy:* For single-molecule experiments, CHO-K1-SNAP_GLP1R cells were seeded onto 25 mm clean glass coverslips at a density of 3x 10^5^ per well. On the following day, cells were labeled in culture medium with 100 pM **LUXendin645** or **LUXendin651** for 20 min. At the end of the incubation, cells were washed 3x 5 min in culture medium. Cells were then imaged at 37 °C in phenol-red free Hank’s balanced salt solution, using a custom built total internal reflection fluorescence microscope (Cairn Research) based on an Eclipse Ti2 (Nikon, Japan) equipped with an EMCCD camera (iXon Ultra, Andor), 637 nm diode laser, and a 100x oil-immersion objective (NA 1.49, Nikon). Image sequences were acquired with an exposure time of 60 ms. Single-molecule image sequences were analyzed with an automated particle detection software (utrack) in the MATLAB environment, as previously described.^53,54^. Data were further analyzed using custom MATLAB algorithms, as previously described.^37,55^

### Two-photon *in vivo* imaging

A 7 week old female C57BL/6J mouse was anesthetized with isoflurane and a small, 1 cm vertical incision was made at the level of the pancreas. The exposed organ was orientated underneath the animal and pressed against a 50 mm glass-bottom dish for imaging on an inverted microscope. Body temperature was maintained using heat pads and heating elements on the objective. The mouse received Hoechst 33342 (1 mg/kg in PBS) to label nuclei, a 150 kDalton fluorescein-conjugated dextran (1 mg/kg in PBS) to label vasculature, and 75 uL of 30 µM **LUXendin555** via retro-orbital IV injection. Images were collected using a Leica SP8 microscope, equipped with a 25x/0.95 NA objective and Spectra Physics MaiTai DeepSee mulitphoton laser. Excitation was delivered at λ = 850 nm, with signals collected with a HyD detector at λ = 460/50, λ = 525/50, λ = 624/40 nm for Hoechst, FITC and **LUXendin555**, respectively. All *in vivo* imaging experiments were performed with approval and oversight from the Indiana University Institutional Animal Care and Use Committee (IACUC).

### Statistical analyses

Measurements were performed on discrete samples unless otherwise stated. All analyses were conducted using GraphPad Prism software. Unpaired or paired Students t-test was used for pairwise comparisons. Multiple interactions were determined using one-way ANOVA followed by Dunn’s or Sidak’s posthoc tests (accounting for degrees of freedom).

### Data availability

The datasets generated during and/or analysed during the current study are available from the corresponding author on reasonable request.

## ACKNOWLEDGMENTS

We thank Bettina Mathes and Alexandra Teslenko for excellent synthetic support. D.J.H. was supported by a Diabetes UK R.D. Lawrence (12/0004431) Fellowship, a Wellcome Trust Institutional Support Award, MRC Confidence in Concept, MRC (MR/N00275X/1) Project and Diabetes UK (17/0005681) Project Grants. ST was supported by an MRC Project Grant (MR/N02589X/1). B.H. was supported by the Wellcome Trust (095101, 200837 and 106130). A.K.L. was supported by R03 DK115990 (to A.K.L.) and Human Islet Research Network UC4 DK104162 (to A.K.L.; RRID:SCR_014393). Intravital microscopy core services were supported by NIH NIDDK Grant P30 DK097512 to the Indiana University School of Medicine. A.T. and B.J. were funded by an MRC Project Grant (MR/R010676/1). D.C. was funded by the Deutsche Forschungsgemeinschaft (SFB/Transregio 166–Project C1) and a Wellcome Trust Senior Research Fellowship. This project has received funding from the European Research Council (ERC) under the European Union’s Horizon 2020 research and innovation programme (Starting Grant 715884 to D.J.H.). We thank Prof Anna Gloyn (University of Oxford) for provision of reagents, Dr Birgit Koch (MPI, Heidelberg) for helpful discussions on *Glp1r^(GE)−/−^* mice and Dr Jacqueline Naylor (MedImmune) for generation of parental SNAP_GLP1R-INS1^GLP1R−/−^ cells.

## CONTRIBUTIONS

J.A., K.J., T.P., J.B. and D.J.H. devised the studies. J.A., A.A., D.N., N.H.F.F., F.B.A., S.T., Z.S., B.H., A.T., T.P., J.B. and D.J.H. performed experiments and analyzed data. J.A. and A.B. generated novel mice. B.J.J. provided reagents. Z.K. and E.D’E. performed super-resolution imaging. C.A.R. and A.K.L. performed *in vivo* imaging experiments. D.C. supervised and analyzed single-molecule microscopy experiments. J.A., K.J., T.P., J.B. and D.J.H. supervised the work. J.A., T.P., J.B. and D.J.H. wrote the manuscript with input from all the authors.

## COMPETING INTERESTS

The authors declare no conflict of interest.

## Notes

#### Summary of Updates

The results section has been updated to clarify GLP1R expression in non-beta cells. Specifically, subtraction of known expression in delta cells, which constitute ~20% insulin-negative cells.

