## Supplementary Information for "LUXendins reveal endogenous glucagon-like peptide-1 receptor distribution and dynamics"

### 1. Chemistry and Spectroscopy

#### 1.1. General

Solvents for chromatography and reactions were purchased HPLC grade (Sigma-Aldrich, 99.8%, extra dry over molecular sieves). If necessary, solvents were degassed either by freeze-pump-thaw or by bubbling $N_2$ through the vigorously stirred solution for several minutes. Unless otherwise stated, all other reagents were used without further purification from commercial sources.

LC-MS was performed on a Shimadzu MS2020 connected to a Nexera UHPLC system equipped with a Waters ACQUITY UPLC BEH C18 (1.7 µm, 50 × 2.1 mm). Buffer A: 0.1% FA in $H_2O$ Buffer B: acetonitrile. The typical gradient was from 10% B for 0.5 min → gradient to 90% B over 4.5 min → 90% B for 0.5 min → gradient to 99% B over 0.5 min with 1 mL/min flow. Retention times ($t_R$) are given in minutes (min).

Preparative and analytical RP-HPLC was performed on a Waters e2695 system equipped with a 2998 PDA detector for product collection (at 220, 550 or 650 nm) on a Supelco Ascentis® C18 HPLC Column (preparative: 5 µm, 250 × 21.2 mm; analytical: 5 µm, 250 × 10 mm). Buffer A: 0.1% TFA in $H_2O$ Buffer B: acetonitrile. The typical gradient was from 10% B for 5 min → gradient to 90% B over 45 min → 90% B for 5 min → gradient to 99% B over 5 min with 8 mL/min flow (preparative) or 4 mL/min (analytical).

NMR spectra were recorded in deuterated solvents on BRUKER Avance III HD 400 (equipped with a CryoProbe™) and calibrated to residual solvent peaks ($^1$H in ppm): DMSO-d6 (2.50/39.52). Multiplicities are abbreviated as follows: s = singlet, d = doublet, t = triplet, q = quartet, br = broad, m = multiplet. Spectra are reported based on appearance, not on theoretical multiplicities derived from structural information.

High resolution mass spectrometry was performed using a Bruker maXis II ETD hyphenated with a Shimadzu Nexera system. The instruments were controlled *via* Brukers otofControl 4.1 and Hystar 4.1 SR2 (4.1.31.1) software. The acquisition rate was set to 3 Hz and the following source parameters were used for positive mode electrospray ionization: End plate offset = 500 V; capillary voltage = 3800 V; nebulizer gas pressure = 45 psi; dry gas flow = 10 L/min; dry temperature = 250 °C. Transfer, quadrupole and collision cell settings are mass range dependent and were fine-adjusted with consideration of the respective analyte's molecular weight. For internal calibration sodium format clusters were used. Samples were desalted *via* fast liquid chromatography. A Supelco Titan™ C18 UHPLC Column, 1.9 µm, 80 Å pore size, 20 × 2.1 mm and a 2 min gradient from 10 to 98% aqueous MeCN with 0.1% FA ($H_2O$: Carl Roth GmbH + Co. KG ROTISOLV® Ultra LC-MS; MeCN: Merck KGaA LiChrosolv® Acetonitrile hypergrade for LC-MS; FA - Merck KGaA LiChropur® Formic acid

98%-100% for LC-MS) was used for separation. Sample dilution in 10% aqueous MeCN (hyper grade) and injection volumes were chosen dependent of the analyte's ionization efficiency. Hence, on-column loadings resulted between 0.25–5.0 ng. Automated internal re-calibration and data analysis of the recorded spectra were performed with Bruker's DataAnalysis 4.4 SR1 software.

UV/Vis spectra were recorded on a Jasco V-770 UV/Vis/NIR Spectrophotometer (PbS-version) with a PAC-743 Peltierthermo 6/8 sample switching unit and a Julabo F-250 cooling system. The spectra were recorded in PBS buffer pH 7.4 using Hellma quartz glass cuvettes (10 mm pathlength).

Fluorescence emission spectra were recorded on a Jasco FP-8600 Fluorescence Spectrometer with PAC-743 Peltierthermo 6/8 sample switching unit and a Julabo F-250 cooling system. The spectra were recorded in PBS buffer pH 7.4 using Hellma dark quartz glass fluorescence cuvettes (10 mm pathlength).

The quantum yields were determined using a Hamamatsu C11347 Absolute PL Quantum Yield Spectrometer (Quantaurus-QY) in PBS pH 7.4.

##### 1.2. General protocol to generate NHS esters

A 1 mL vial was charged with 1.0 equiv. of acid dissolved in DMF (1 mL / 10 mg) and 4.0 equiv. of DIPEA was added before 1.1 equiv. of TSTU in one portion (for amounts <1 mg of TSTU, stock solutions were prepared as it is critical to not overload TSTU). The active NHS ester was allowed to form for 15 min and used without further purification.

##### 1.3. General protocol for peptide coupling using NHS esters

A 1 mL vial was charged with 1.0 equiv. amine dissolved in DMF (1 mL / 10 mg) and 4.0 equiv. DIPEA. The pre-formed NHS ester (section 1.3) was added drop-wise at and the reaction mixture was allowed to stir at r.t. Upon complete conversion according to LCMS (usually <30 min), the reaction was quenched by addition of 5 vol% HOAc and 10 vol% water and subjected to RP-HPLC. The desired products were obtained as colored powders.

##### 1.4. TMR-Mal

Red powder.

$^{1}$**H NMR** (400 MHz, DMSO-$d_6$): $\delta$ [ppm] = 8.78 (t, J = 6.0 Hz, 1H), 8.08–8.02 (m, 2H), 7.51 (d, J = 1.3 Hz, 1H), 6.95 (s, 2H), 6.51 (d, J = 1.4 Hz, 6H), 3.52 (t, J = 5.8 Hz, 2H), 3.33 (2H), 2.94 (s, 12H). The signal at 3.33 ppm is colocated with the water signal at 3.31 ppm, but was detectable via COSY and HSQC spectroscopy.

**$^{13}$C NMR** (100 MHz, DMSO-*$d_6$*): $\delta$ [ppm] = 171.01, 168.24, 165.04, 152.85, 152.14, 151.97, 140.46, 134.53, 128.99, 128.59, 128.48, 124.76, 122.29, 109.09, 105.59, 97.98, 84.74, 37.81, 36.91.

**UV/Vis** (LCMS): $\lambda_{max}$ = 556 nm.

***$t_R$*** (LCMS) = 2.502 min.

**HRMS** (ESI): calc. for $C_{31}H_{29}N_4O_6$ $[M+H]^+$: 553.2082, found: 553.2083.

#### 1.5. Cy5-Mal

Blue powder.

**$^1$H NMR** (400 MHz, MeOH-*$d_4$*): $\delta$ [ppm] = 8.51 (s, 1H), 8.24 (t, J = 13.3 Hz, 2H), 7.49 (d, J = 7.4 Hz, 2H), 7.42 (t, J = 7.8 Hz, 2H), 7.28 (dt, J = 15.5, 7.2 Hz, 4H), 7.06 (dt, J = 29.3, 9.8 Hz, 2H), 6.82 – 6.75 (m, 2H), 6.67 – 6.53 (m, 2H), 6.28 (t, J = 13.9 Hz, 2H), 4.11 (t, J = 7.5 Hz, 2H), 3.66 – 3.55 (m, 6H), 2.14 (t, J = 7.3 Hz, 2H), 1.81 (t, J = 7.7 Hz, 2H), 1.65 (t, J = 7.7 Hz, 2H), 1.55 (s, 3H), 1.46 (d, J = 7.4 Hz, 4H), 1.36 – 1.19 (m, 2H), 1.03 (s, 1H).

**UV/Vis** (LCMS): $\lambda_{max}$ = 640 nm.

***$t_R$*** (LCMS) = 2.971 min.

**HRMS** (ESI): calc. for $C_{38}H_{45}N_4O_3$ $[M]^+$: 605.3486, found: 605.3488.

#### 1.6. SiR-Mal

Blue powder.

**$^1$H NMR** (400 MHz, DMSO-*$d_6$*): $\delta$ [ppm] = 8.82 (s, 1H), 8.02 (d, J = 8.0 Hz, 1H), 7.62 (s, 1H), 7.57 (d, J = 1.3 Hz, 1H), 7.02 (t, J = 2.7 Hz, 3H), 6.96 (s, 2H), 6.78 – 6.49 (m, 5H), 3.33 (2H), 2.92 (d, J = 1.4 Hz, 12H), 0.64 (d, J = 2.3 Hz, 3H), 0.53 (d, J = 3.7 Hz, 3H). The signal at 3.33 ppm is colocated with the water signal at 3.31 ppm, but was detectable via COSY and HSQC spectroscopy.

**$^{13}$C NMR** (100 MHz, DMSO-*$d_6$*): $\delta$ [ppm] = 171.02, 169.26, 165.25, 154.68, 149.25, 149.18, 139.88, 135.85, 135.29, 134.51, 130.43, 128.00, 127.34, 127.26, 125.41, 122.72, 116.48, 116.41, 113.68, 91.26, 40.20, 40.15, 39.99, 39.94, 39.73, 39.52, 39.31, 39.11, 38.89, 37.83, 36.90, 0.05, -1.24.

**UV/Vis** (LCMS): $\lambda_{max}$ = 656 nm.

***$t_R$*** (LCMS) = 2.514 min.

**HRMS** (ESI): calc. for $C_{33}H_{35}N_4O_5Si$ $[M+H]^+$: 595.2371, found: 595.2374.

**1.7. Synthesis of S39C-Ex4(9-39)**

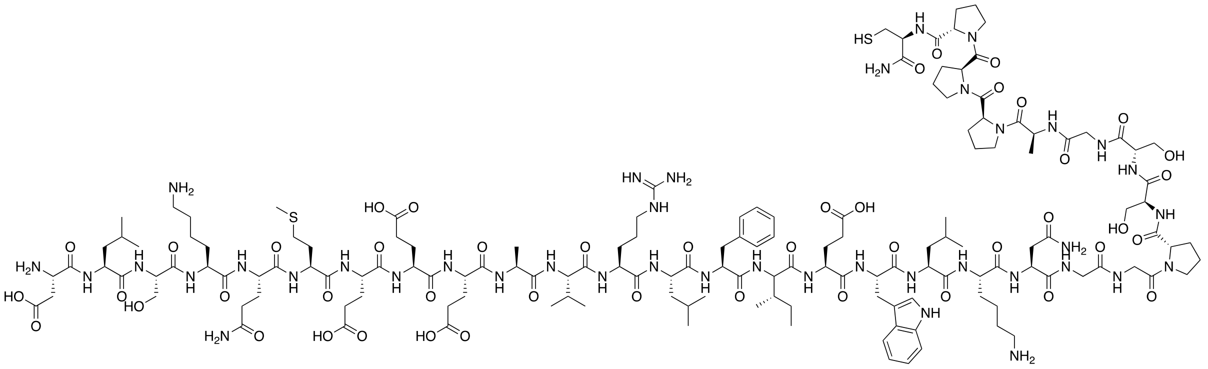

**S39C-Ex4(9-39)**

H-Asp-Leu-Ser-Lys-Gln-Met-Glu-Glu-Glu-Ala-Val-Arg-Leu-Phe-Ile-Glu-Trp-Leu-Lys-Asn-Gly-Gly-Pro-Ser-Ser-Gly-Ala-Pro-Pro-Pro-Cys-$NH_2$

Chemical Formula: $C_{149}H_{234}N_{40}O_{46}S_2$
Exact Mass: 3383,6642
Molecular Weight: 3385,8650

Peptides were synthesized on a CEM Liberty Blue Peptide Synthesizer with a CEM Discovery Microwave using standard Fmoc-protected solid phase peptide synthesis protocols with standard reagents. Pre-loaded Fmoc-Cys(Trt)-Tentagel S PHB resin (Rapp Polymere, Germany) containing 0.2-0.3 mmol/g amino acid was used as solid-phase. Peptide synthesis scale was 0.1 mmol using the standard coupling reagents DIC/Oxyma 0.5/1.0 M in DMF and DIPEA 2 M in DMF. Fmoc-protected amino acids (Sigma-Aldrich and NovaBiochem Merck, Germany) with standard residual protecting groups were coupled using a five-fold excess (2 M solutions). Deprotection of the Fmoc-protecting group was achieved by treatment with 20% piperidine in DMF. After completion of all coupling steps, the resin-bound peptide was transferred into a syringe with frit followed by global deprotection using 10 mL of a TFA:$H_2O$:tri-*iso*-propylsilane (95:2.5:2.5) mixture within 2 h under argon atmosphere. The peptide solution was filtered and the filtrate was concentrated under reduced pressure and residual TFA was removed by co-evaporation with toluene (3x). The residue was dissolved in a small amount of methanol and precipitated in 40 mL chilled diethyl ether and stored overnight at –38 °C to complete precipitation. The suspension was subjected to centrifugation (3200 rpm, 4 °C, 5 min), the supernatant removed and the residue dried before being reconstituted in water and subjected to RP-HPLC purification. The combined purified fractions were lyophilized and the **S39C-Ex4(9-39)** peptide was yielded as white TFA salt.

**UV/Vis** (LCMS): $\lambda_{max}$ = 212 nm.

$\boldsymbol{t_R}$ (HPLC) = 24.4 min.

**HRMS** (ESI): calc. for $C_{149}H_{238}N_{40}O_{46}S_2$ $[M+4H]^{4+}$: 847.1742, found: 847.1741.

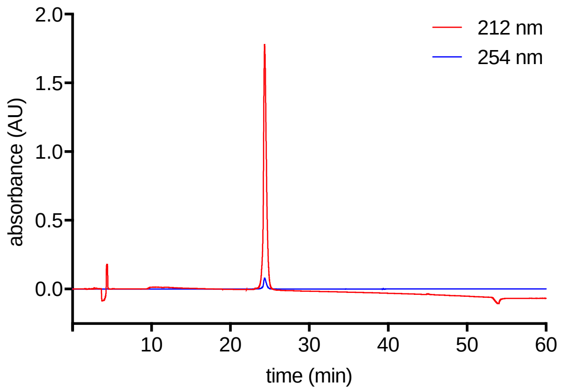


Analytical RP-HPLC trace of **S39C-Ex4(9-39)** peptide.

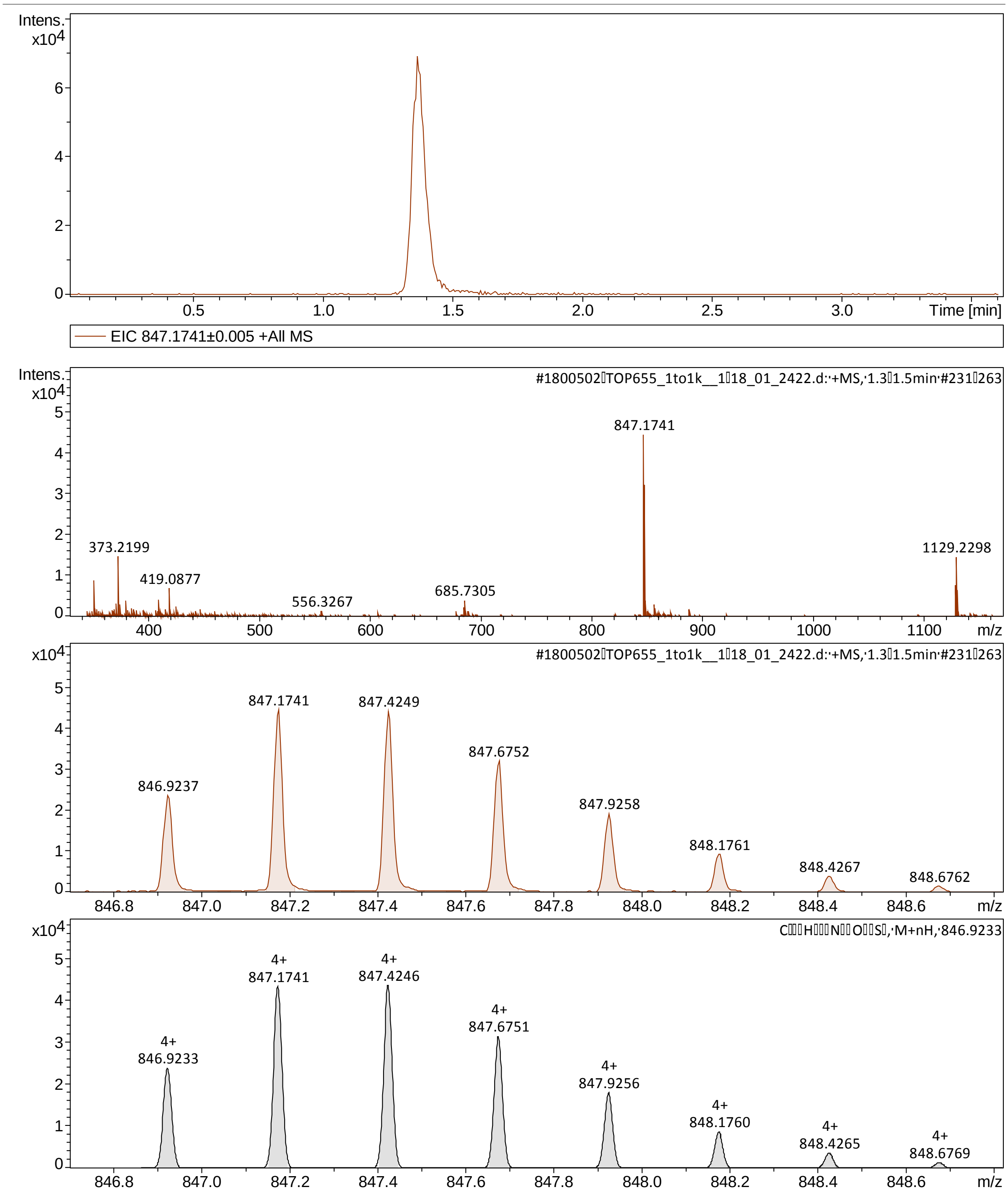


HR ESI-MS spectrum of S39C-Ex4(9-39).

#### 1.8. Synthesis of LUXendin555

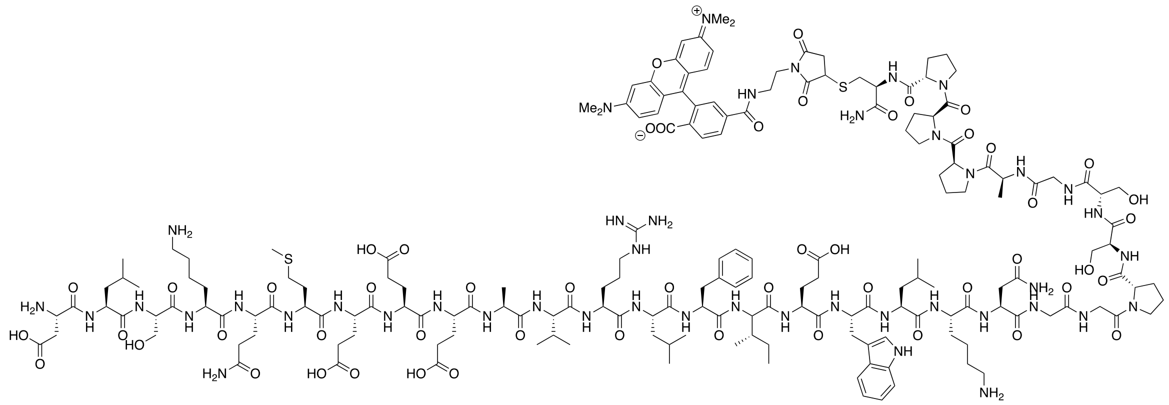

**LUXendin550**

H-Asp-Leu-Ser-Lys-Gln-Met-Glu-Glu-Glu-Ala-Val-Arg-Leu-Phe-Ile-Glu-Trp-Leu-Lys-Asn-Gly-Gly-Pro-Ser-Ser-Gly-Ala-Pro-Pro-Pro-Cys(TMR)-$NH_2$

Chemical Formula: $C_{180}H_{262}N_{44}O_{52}S_2$
Exact Mass: 3935,8651
Molecular Weight: 3938,4520

To a solution of S39C-Ex4(9-39) (2.03 mg, 600 nmol, 1.0 eq.) in PBS (400 uL) was added TMR-Mal (0.5 mg, 877 nmol, 1.5 eq.). The solution was stirred at room temperature over night before being subjected to RP-HPLC purification (water/ACN gradient, 90/10 → 10/90 in 60 min). The purified fractions were combined and lyophilized from the HPLC solvent to yield the **LUXendin555** as light blue TFA salt. The concentration was determined *via* the absorption of the TMR fluorophore at 550 nm (ε = 80000 mol $L^{-1}$ $cm^{-1}$ in PBS with 0.1% SDS) and stocks of 10 nmol each were prepared and stored at -80 °C.

**UV/Vis** (UV, FluoSpec): $Ex_{max}$ = 555 nm, $Em_{max}$ = 579.
$\boldsymbol{t_R}$ (HPLC) = 25.0 min.
**HRMS** (ESI): calc. for $C_{180}H_{266}N_{44}O_{52}S_2$ $[M+4H]^{4+}$: 985.4764, found: 985.4749.

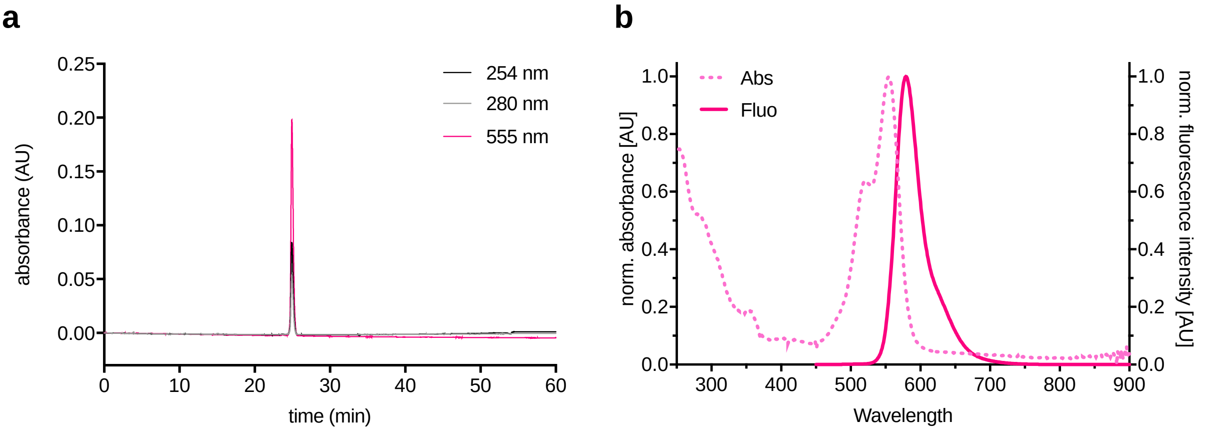


(**a**) Analytical RP-HPLC trace of **LUXendin555**. (**b**) Normalized absorption and fluorescence emission spectra of **LUXendin555**.

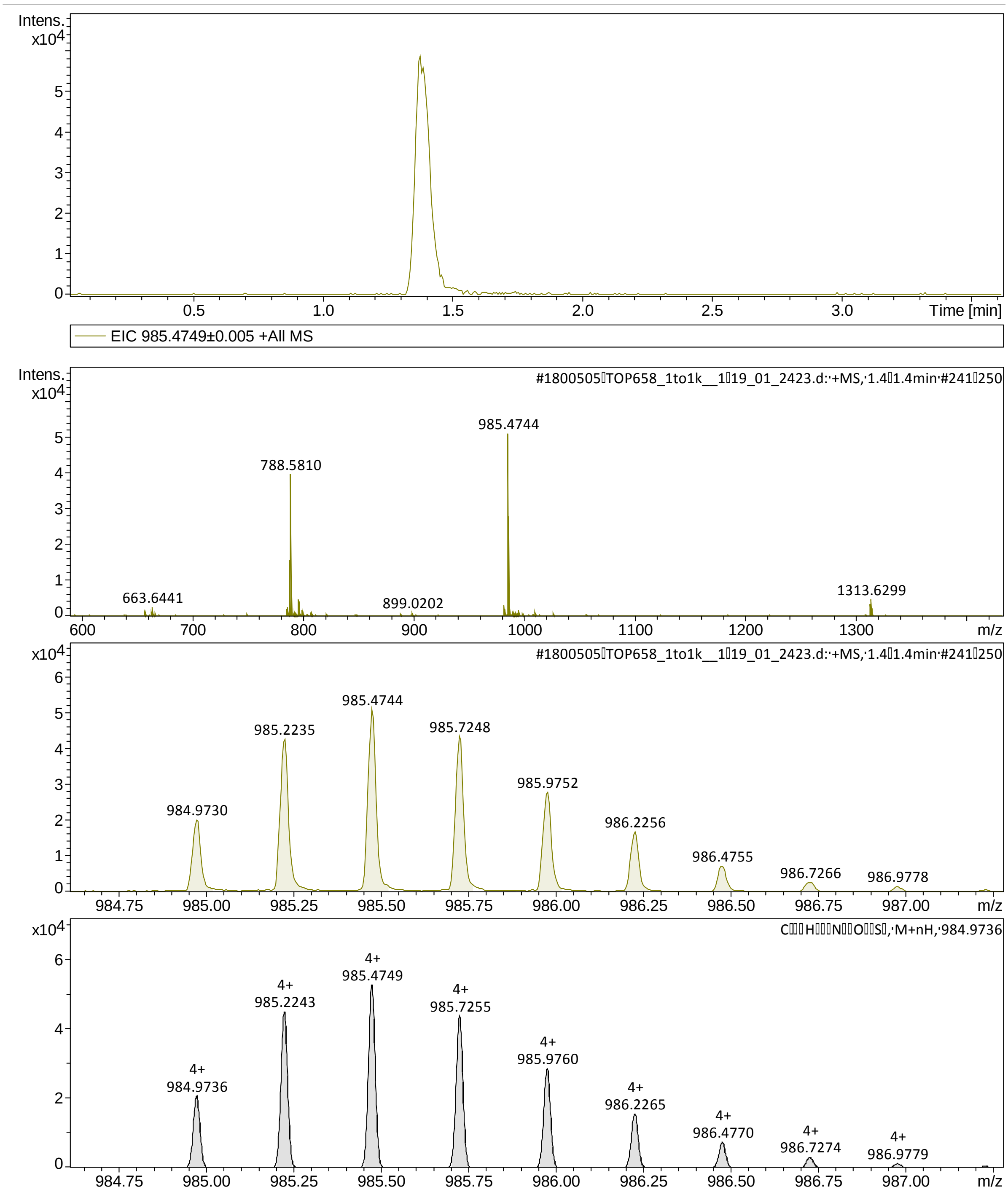


HR ESI-MS of LUXendin555.

#### 1.9. Synthesis of LUXendin645

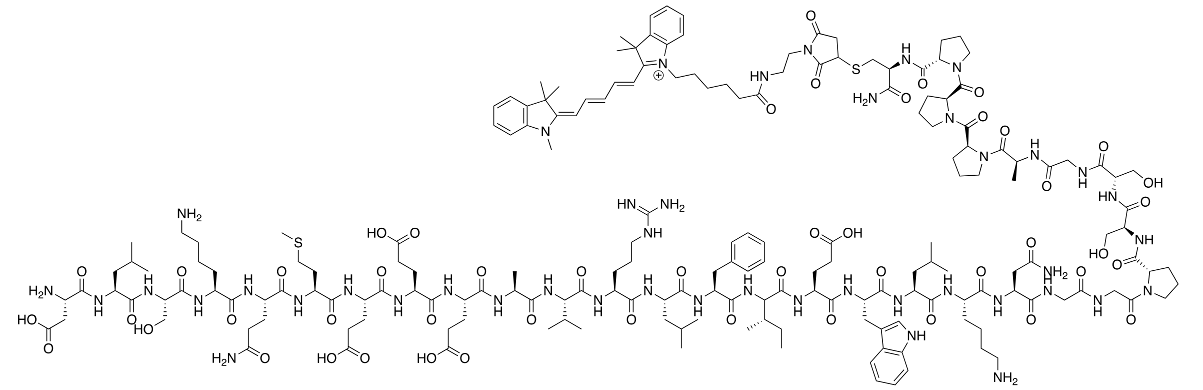

**LUXendin645**

$H\text{-}Asp\text{-}Leu\text{-}Ser\text{-}Lys\text{-}Gln\text{-}Met\text{-}Glu\text{-}Glu\text{-}Glu\text{-}Ala\text{-}Val\text{-}Arg\text{-}Leu\text{-}Phe\text{-}Ile\text{-}Glu\text{-}Trp\text{-}Leu\text{-}Lys\text{-}Asn\text{-}Gly\text{-}Gly\text{-}Pro\text{-}Ser\text{-}Ser\text{-}Gly\text{-}Ala\text{-}Pro\text{-}Pro\text{-}Pro\text{-}Cys(Cy5)\text{-}NH_2$

Chemical Formula: $C_{187}H_{279}N_{44}O_{49}S_2^{+}$
Exact Mass: 3989,0128
Molecular Weight: 3991,6675

To a solution of S39C-Ex4(9-39) (2.03 mg, 600 nmol, 1.0 eq.) in PBS (400 uL) was added Cy5-Mal (0.5 mg, 824 nmol, 1.4 eq.). The solution was stirred at room temperature over night before being subjected to RP-HPLC purification (water/ACN gradient, 90/10 → 10/90 in 60 min). The purified fractions were combined and lyophilized from the HPLC solvent to yield the **LUXendin645** as light blue TFA salt. The concentration was determined *via* the absorption of the Cy5 fluorophore at 647 nm ($\varepsilon$ = 250000 mol $L^{-1}$ $cm^{-1}$ in PBS with 0.1% SDS) and stocks of 10 nmol each were prepared and stored at -80 °C.

**UV/Vis** (UV, FluoSpec): $Ex_{max}$ = 645 nm, $Em_{max}$ = 664.

$\boldsymbol{t_R}$ (HPLC) = 26.3 min.

**HRMS** (ESI): calc. for $C_{187}H_{279}N_{44}O_{49}S_2$ $[M+4H]^{5+}$: 799.0097, found: 799.0095.

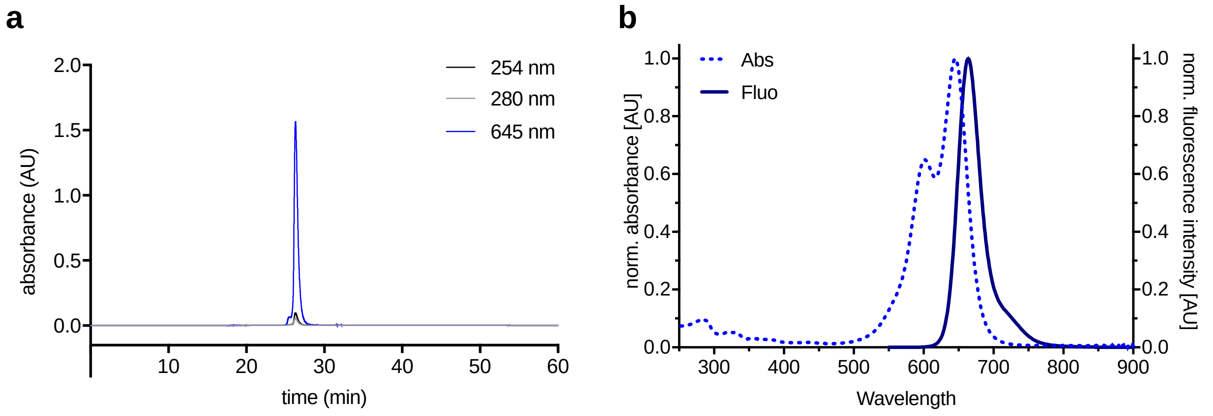


(**a**) Analytical RP-HPLC trace of **LUXendin645**. (**b**) Normalized absorption and fluorescence emission spectra of **LUXendin645**.

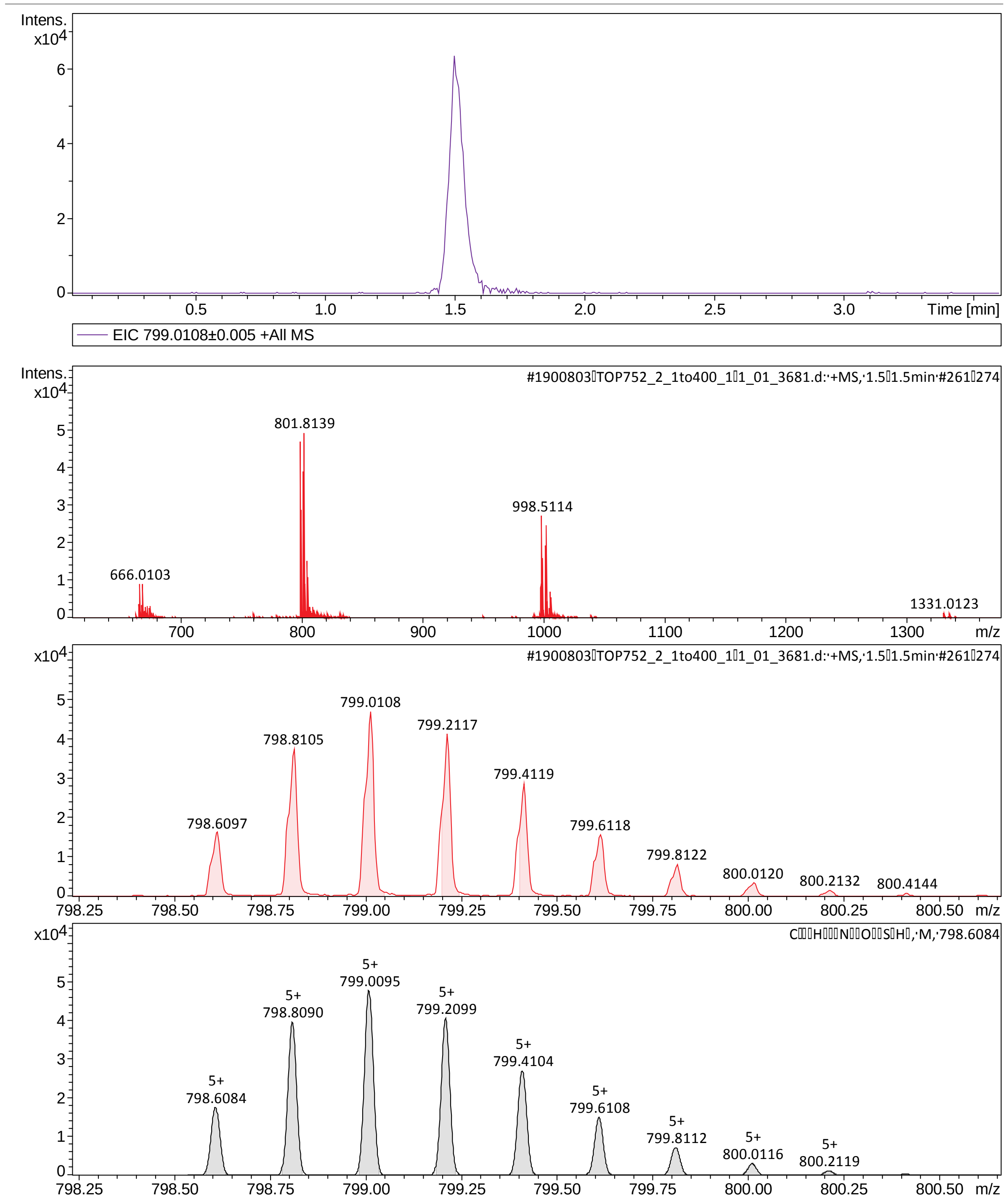


HR ESI-MS of LUXendin645.

#### 1.10. Synthesis of LUXendin651

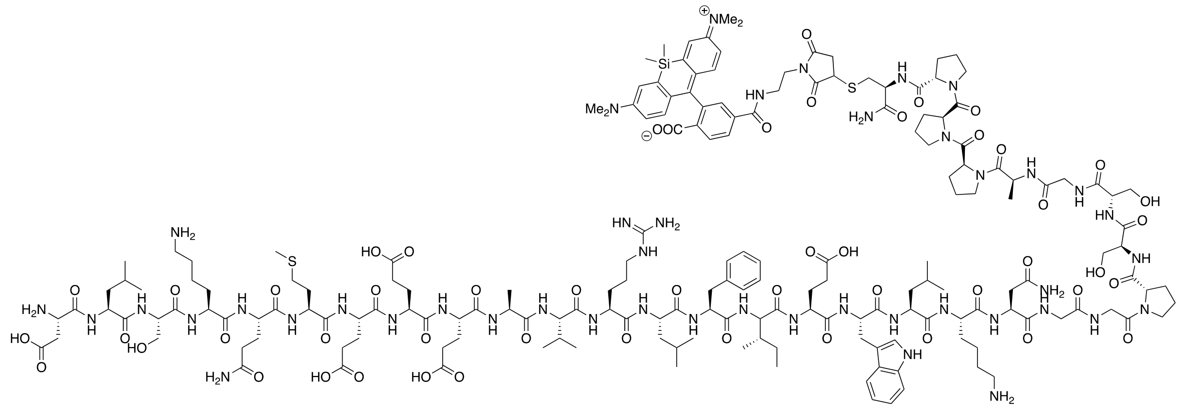

**LUXendin651**

H-Asp-Leu-Ser-Lys-Gln-Met-Glu-Glu-Glu-Ala-Val-Arg-Leu-Phe-Ile-Glu-Trp-Leu-Lys-Asn-Gly-Gly-Pro-Ser-Ser-Gly-Ala-Pro-Pro-Pro-Cys(SiR)-$NH_2$

Chemical Formula: $C_{182}H_{268}N_{44}O_{51}S_2Si$
Exact Mass: 3977,8941
Molecular Weight: 3980,6080

To a solution of S39C-Ex4(9-39) (2.03 mg, 600 nmol, 1.0 eq.) in PBS (400 uL) was added SiR-Mal (0.5 mg, 833 nmol, 1.4 eq.). The solution was stirred at room temperature over night before being subjected to RP-HPLC purification (water/ACN gradient, 90/10 → 10/90 in 60 min). The purified fractions were combined and lyophilized from the HPLC solvent to yield the **LUXendin651** as light blue TFA salt. The concentration was determined *via* the absorption of the SiR fluorophore at 647 nm (ε = 100000 mol $L^{-1}$ $cm^{-1}$ in PBS with 0.1% SDS) and stocks of 10 nmol each were prepared and stored at -80 °C.

**UV/Vis** (UV, FluoSpec): $Ex_{max}$ = 651 nm, $Em_{max}$ = 669.

***$t_R$*** (HPLC) = 24.8 min.

**HRMS** (ESI): calc. for $C_{182}H_{269}N_{44}O_{51}S_2Si$ $[M+4H]^{4+}$: 995.9836, found: 995.9821.

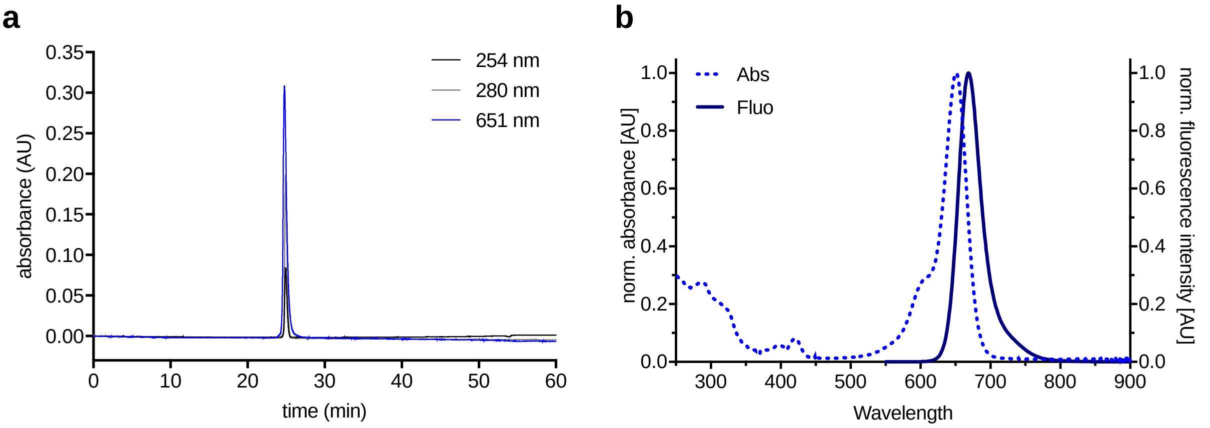


(**a**) Analytical RP-HPLC trace of **LUXendin651**. (**b**) Normalized absorption and fluorescence emission spectra of **LUXendin651**.

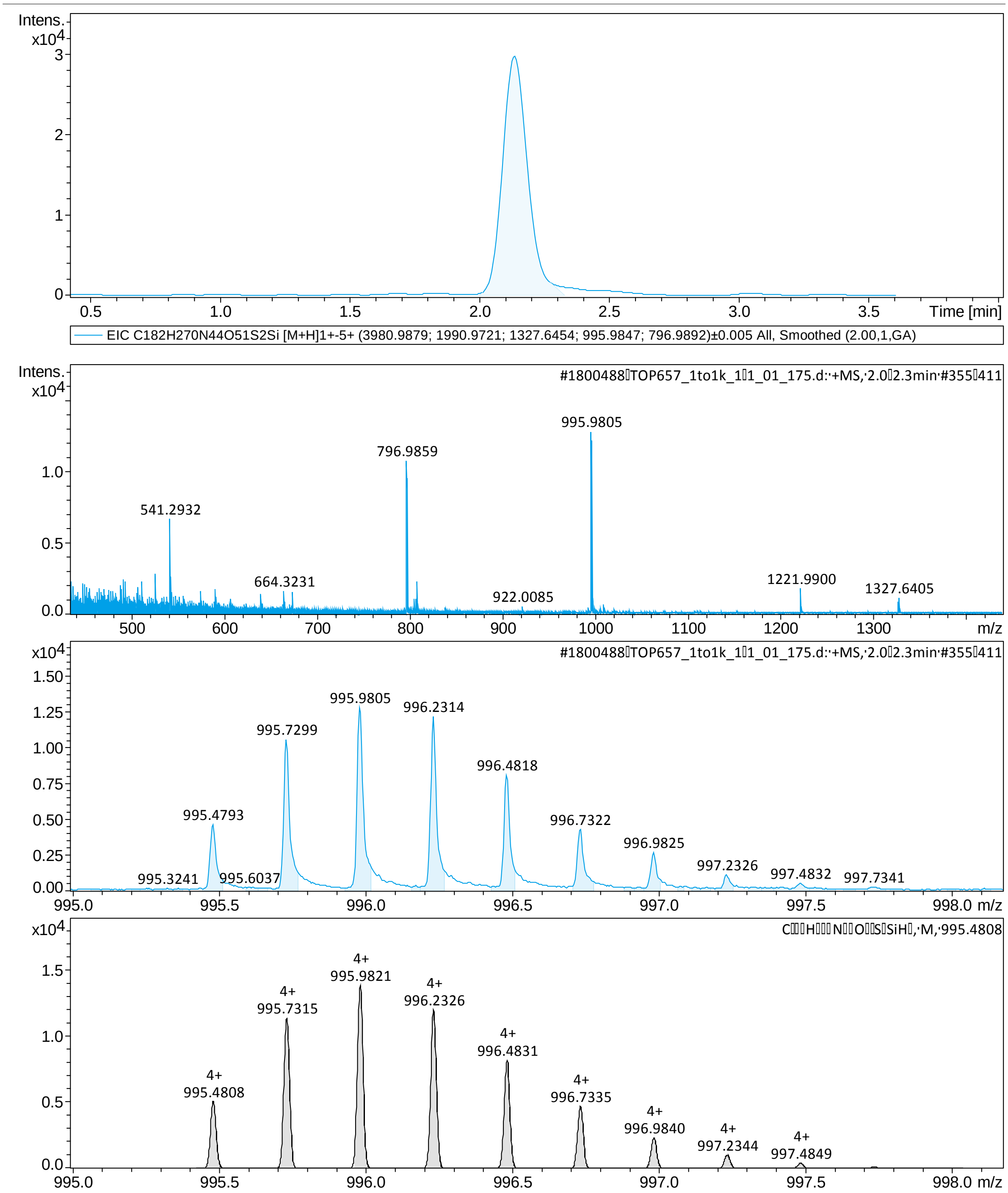


HR ESI-MS of LUXendin651.

**SUPPLEMENTARY FIGURES**

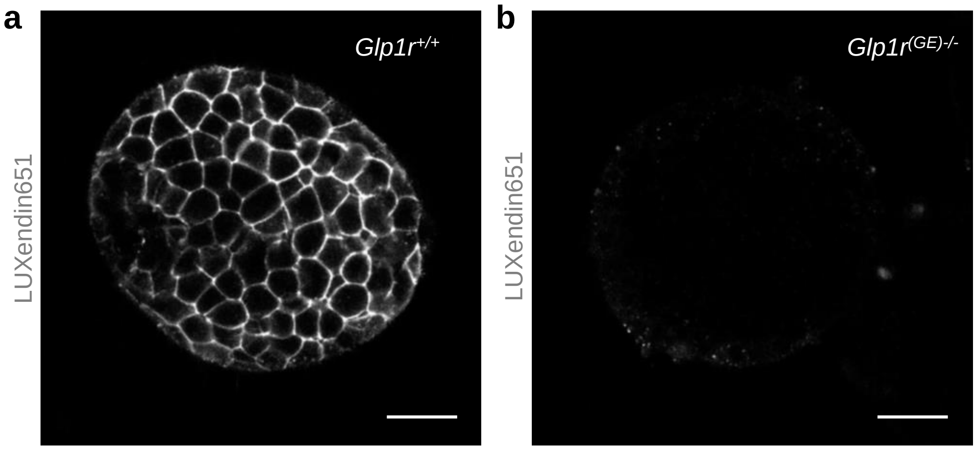


**Figure S1: LUXendin651 does not label islets from *Glp1r*$^{(GE)-/-}$ animals. a** and **b**, Signal can be detected in wild-type (**a**) but not *Glp1r*$^{(GE)-/-}$ islets (**b**) (n = 6 islets) (scale bar = 26.5 µm).

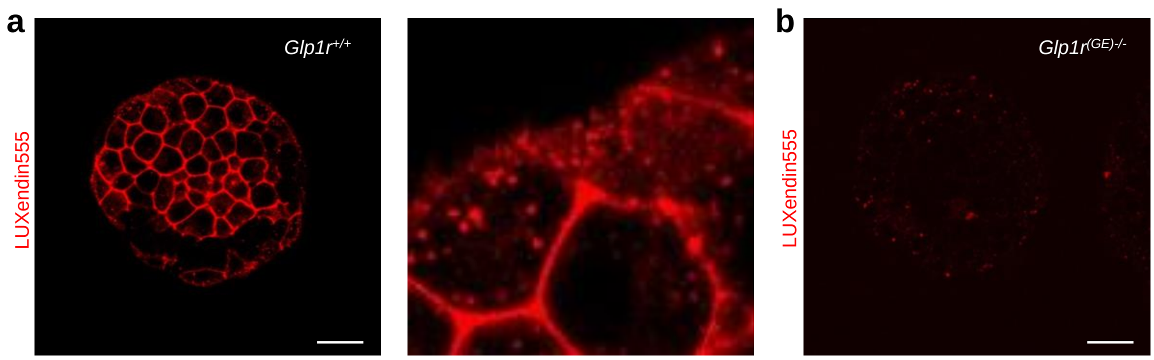


**Figure S2: LUXendin555 does not label islets from *Glp1r*$^{(GE)-/-}$ animals. a** and **b**, Signal can be detected in wild-type (**a**) but not *Glp1r*$^{(GE)-/-}$ islets (**b**) labelled with 100 nM **LUXendin555** (inset shows internalized GLP1R) (n = 6 islets) (scale bar = 26.5 µm). Note that the brightness/contrast in image (**b**) has been increased relative to image (**a**) to allow the *Glp1r*$^{(GE)-/-}$ islet to be seen.

#### SUPPLEMENTARY MOVIE LEGENDS

**Movie S1:** Two-photon z-stack of **LUXendin645**-labeled islets (147 µm).

**Movie S2:** Single-molecule localization microscopy in **LUXendin645**-labeled CHO-K1-SNAP_GLP1R cells (95 nm per pixel) (20 frames per second).

**Movie S3:** Single-molecule localization microscopy in **LUXendin651**-labeled CHO-K1-SNAP_GLP1R cells (95 nm per pixel) (20 frames per second).

**Movie S4:** Single particle tracking in **LUXendin651**-labeled CHO-K1-SNAP_GLP1R cells (47.5 nm per pixel) (20 frames per second).
